# Characterization of Pch2 localization determinants reveals a nucleolar-independent role in the meiotic recombination checkpoint

**DOI:** 10.1101/541367

**Authors:** Esther Herruzo, Beatriz Santos, Raimundo Freire, Jesús A. Carballo, Pedro A. San-Segundo

**Affiliations:** Instituto de Biología Funcional y Genómica (IBFG). Consejo Superior de Investigaciones Científicas (CSIC) and University of Salamanca. 37007-Salamanca, Spain; Departamento de Microbiología y Genética. University of Salamanca. 37007-Salamanca, Spain; Hospital Universitario de Canarias, Instituto de Tecnologías Biomédicas. 38320-La Laguna, Tenerife, Spain; Department of Cellular and Molecular Biology. Centro de Investigaciones Biológicas. Consejo Superior de Investigaciones Científicas (CSIC). 28040-Madrid, Spain

**Keywords:** meiosis, checkpoint, synapsis, recombination, Pch2, Orc1

## Abstract

The meiotic recombination checkpoint blocks meiotic cell cycle progression in response to synapsis and/or recombination defects to prevent aberrant chromosome segregation. The evolutionarily-conserved budding yeast Pch2^TRIP13^ AAA+ ATPase participates in this pathway by supporting phosphorylation of the Hop1^HORMAD^ adaptor at T318. In the wild type, Pch2 localizes to synapsed chromosomes and to the unsynapsed rDNA region (nucleolus), excluding Hop1. In contrast, in synaptonemal complex (SC)-defective *zip1Δ* mutants, which undergo checkpoint activation, Pch2 is detected only on the nucleolus. Alterations in some epigenetic marks that lead to Pch2 dispersion from the nucleolus suppress *zip1Δ*-induced checkpoint arrest. These observations have led to the notion that Pch2 nucleolar localization could be important for the meiotic recombination checkpoint. Here we investigate how Pch2 chromosomal distribution impacts on checkpoint function. We have generated and characterized several mutations that alter Pch2 localization pattern resulting in aberrant Hop1 distribution and compromised meiotic checkpoint response. Besides the AAA+ signature, we have identified a basic motif in the extended N-terminal domain critical for Pch2’s checkpoint function and localization. We have also examined the functional relevance of the described Orc1-Pch2 interaction. Both proteins colocalize in the rDNA, and Orc1 depletion during meiotic prophase prevents Pch2 targeting to the rDNA allowing unwanted Hop1 accumulation on this region. However, Pch2 association with SC components remains intact in the absence of Orc1. We finally show that checkpoint activation is not affected by the lack of Orc1 demonstrating that, in contrast to previous hypotheses, nucleolar localization of Pch2 is actually dispensable for the meiotic checkpoint.

## INTRODUCTION

During gametogenesis, a tight spatiotemporal control of a myriad of interrelated events that integrate the meiotic program must occur in order to achieve the successful generation of gametes with the adequate chromosome complement. This control is reinforced by the action of surveillance mechanisms, or checkpoints, that block meiotic progression in response to defects in critical meiotic processes thus preventing errors in the distribution of chromosomes to the meiotic progeny (Subramanian and Hochwagen 2014). Checkpoint pathways involve a series of molecular events frequently relying on protein phosphorylation, to eventually give rise to the adequate cellular responses including cell-cycle arrest among others. The so-called pachytene checkpoint or meiotic recombination checkpoint operates during meiosis to face failures in the synapsis and/or recombination processes. Depending on the nature of the triggering signal, different sensing mechanisms are involved. For example, while RPA-coated processed meiotic DNA double-strand breaks (DSBs) activate the Mec1^ATR^ sensor kinase via 9-1-1 complex and Ddc2^ATRIP^-mediated recruitment (Lydall et al. 1996; Hong and Roeder 2002; Eichinger and Jentsch 2010; Refolio et al. 2011), unresected DSBs activate Tel1^ATM^ via the Mre11-Rad50-Xrs2^NBS1^ (MRX) complex (Usui et al. 2001). In any case, irrespective of the checkpoint inducing event the final outcome involves a block in meiotic progression by down-regulation of the cell cycle machinery (Acosta et al. 2011; Prugar et al. 2017).

The evolutionarily conserved Pch2^TRIP13^ protein was initially discovered in *Saccharomyces cerevisiae* in a genetic screen for mutations that alleviate the checkpoint-induced meiotic arrest of the *zip1Δ* mutant lacking a main component of the central region of the synaptonemal complex (SC) (Sym et al. 1993; San-Segundo and Roeder 1999; Wu and Burgess 2006; Herruzo et al. 2016). Pch2 is also required for the checkpoint response elicited by unresected DSBs involving the interaction with Xrs2 (Ho and Burgess 2011). The participation of Pch2 orthologs in the checkpoint response to various meiotic stimuli has been also reported in other organisms, such as for example worms (Bhalla and Dernburg 2005) and flies (Joyce and McKim 2009). Besides the checkpoint role, Pch2 additionally impinges on multiple interrelated meiotic recombination events, including DSB formation (Farmer et al. 2012; Joshi et al. 2015), chromosome axis morphogenesis (Börner et al. 2008; Joshi et al. 2009), crossover control (Medhi et al. 2016; Chakraborty et al. 2017), interhomolog bias (Zanders et al. 2011; Subramanian et al. 2016), crossover interference (Zanders and Alani 2009) and ribosomal DNA (rDNA) array stability (San-Segundo and Roeder 1999; Vader et al. 2011). Pch2^TRIP13^ belongs to the AAA+ family of ATPases (Chen et al. 2014; Vader 2015) that utilize the energy generated from ATP hydrolysis to produce conformational changes on the substrates (Hanson and Whiteheart 2005). In the case of Pch2^TRIP13^ many of its meiotic functions involve the action on the Hop1^HORMAD1,2^ SC component; in particular, Pch2 promotes Hop1 disengagement from chromosome axes as synapsis progresses (San-Segundo and Roeder 1999; Li and Schimenti 2007; Börner et al. 2008; Wojtasz et al. 2009; Roig et al. 2010; Herruzo et al. 2016; Subramanian et al. 2016). Although budding yeast *PCH2* is only expressed in meiotic cells, recent studies have revealed a crucial role for Pch2 orthologs in the mitotic spindle assembly checkpoint in worms and mammals, also acting on a HORMA-domain containing protein, namely MAD2 (Nelson et al. 2015; Ye et al. 2015; Ma and Poon 2018; West et al. 2018).

Since Pch2 removes Hop1 from meiotic chromosomes, the *pch2*Δ single mutant displays more abundant and continuous Hop1 distribution on synapsed chromosomes (San-Segundo and Roeder 1999; Börner et al. 2008). In contrast, unexpectedly, our previous work has demonstrated that under checkpoint-inducing conditions (*zip1Δ*), the Pch2 protein is critically required for maintaining linear Hop1 localization along chromosome axes and, more important, for sustaining high levels of Hop1 phosphorylation at Thr318. In other words, chromosomal Hop1 is less abundant and Hop1-T318 phosphorylation is drastically reduced in *zip1Δ pch2*Δ compared to *zip1Δ* (Herruzo et al. 2016). Deficient Mec1-dependent Hop1-T318 phosphorylation leads to impaired Mek1 activation (Carballo et al. 2008), thus explaining the defective checkpoint response in *zip1*Δ *pch2*Δ cells. Importantly, *HOP1* overexpression restores checkpoint function in *zip1*Δ *pch2*Δ (Herruzo et al. 2016). Cytological studies have uncovered a peculiar localization pattern for the Pch2 protein on meiotic chromosomes. Pch2 displays a prominent localization in the unsynapsed rDNA region of chromosome XII and a weaker distribution on interstitial synapsed chromosomal sites (San-Segundo and Roeder 1999; Börner et al. 2008; Herruzo et al. 2016). The association of Pch2 with the SC is clearly evidenced by the presence of Pch2 on Zip1-containing polycomplexes (San-Segundo and Roeder 1999; Dong and Roeder 2000). Remarkably, in a checkpoint-activated scenario like the SC-deficient *zip1Δ* mutant, Pch2 has been only detected in the nucleolar region. Nucleolar accumulation of Pch2 requires histone H3K79 methylation by Dot1 and proper levels of H4K16 acetylation controlled by Sir2 (Ontoso et al. 2013; Cavero et al. 2016). The fact that both *dot1* and *sir2* mutations impair the meiotic recombination checkpoint is consistent with the notion that nucleolar Pch2 is important for checkpoint activity. However, this hypothesis has not yet been tested directly.

In this work, we have identified a basic motif in the non-conserved N-terminal domain (NTD) of Pch2 that is necessary for its localization to both SC and rDNA. Mutation of this motif results in impaired checkpoint response suggesting that it may be required for proper Pch2 chromatin association and/or interaction with additional critical factors. The Orc1 protein targets Pch2 to the rDNA to repress meiotic DSB formation (Vader et al. 2011); thus, in order to definitely assess the functional relevance of Pch2 nucleolar localization for the *zip1Δ* -induced checkpoint we have engineered a conditional *orc1-3mAID* degron allele. We found that induced Orc1 degradation during meiotic prophase precludes Pch2 localization to the rDNA, but association with SC components is unaltered. Using various cytological and molecular assays, we show that checkpoint activation remains intact in the absence of Orc1. Thus, we conclude that Pch2 nucleolar localization is dispensable for the checkpoint response to SC defects.

## RESULTS

### An NLS-like element in the Pch2 N-terminal domain is required for checkpoint function and localization

Alignment of the proteins sequences of Pch2/TRIP13 orthologs of different species revealed the presence of a non-conserved extended N-terminal domain (NTD) in the Pch2 protein of *S*. *cerevisiae* (Fig. 1a and Fig. S1). In wild-type yeast meiotic chromosomes Pch2 accumulates at the SC-devoid nucleolar rDNA region of chromosome XII and a minor fraction also associates to the SC along synapsed chromosomes (San-Segundo and Roeder 1999; Börner et al. 2008; Herruzo et al. 2016; Subramanian et al. 2016). In other organisms such as, for example, plants and worms, Pch2 orthologs have been localized only to the SC (Miao et al. 2013; Deshong et al. 2014; Lambing et al. 2015). The fact that nucleolar accumulation of Pch2 appears to be restricted to budding yeast suggests that the NTD of Pch2 may be involved in this characteristic distribution pattern. Several observations point to a critical role for the nucleolar Pch2 in meiotic recombination checkpoint function (see Introduction); therefore, we searched the NTD sequence for motifs possibly involved in nucleolar targeting. We found a 17-amino-acid stretch at positions 42 to 58 containing several basic residues that could resemble a nuclear or nucleolar localization signal (NLS/NoLS) (Fig. 1a and Fig. S1). To explore the meiotic relevance of this NLS-like sequence, we used the *delitto perfetto* approach to generate *PCH2* mutants carrying a precise deletion of this motif in the genomic loci (*pch2-nlsΔ*) (Fig. 1a). Like the *pch2Δ* null mutant, the *pch2-nlsΔ* single mutant completed meiosis and sporulation with similar kinetics and efficiency than the wild type generating high levels of viable spores (Fig. 1b, c; Table 1). Notably, when the checkpoint was triggered by the absence of Zip1, the *pch2-nlsΔ* mutation suppressed the sporulation defect of *zip1Δ* (Fig. 1b) to produce largely inviable spores (Table 1). Likewise, the substantial delay in the kinetics of meiotic divisions of *zip1Δ* was drastically suppressed in *zip1Δ pch2-nlsΔ* to reach near wild-type kinetics (Fig. 1d). Western blot analysis revealed that, in both the wild-type strain and the *pch2-nlsΔ* single mutant, the Pch2 and Pch2-nlsΔ proteins were induced during meiotic prophase (15 h) and then disappeared with similar kinetics as meiosis and sporulation progresses (Herruzo et al. 2016) (Fig. 1c, e). In the *zip1Δ* mutant, high levels of the wild-type Pch2 protein persisted until late time points (Fig. 1e) according with its strong meiotic prophase block (Fig. 1d). In contrast, in *zip1Δ pch2-nlsΔ*, the levels of Pch2-nlsΔ drastically diminished as meiotic divisions took place (Fig. 1d,e). To determine whether the reduced levels of Pch2-nlsΔ in *zip1Δ pch2-nlsΔ* cells were responsible for the bypass of *zip1Δ* arrest or simply reflected the consequences of meiotic progression beyond the point when the Pch2 protein is normally produced, we quantified Pch2 protein levels in the *ndt80Δ* mutant at the 24-hour time point, when most cells in the BR strain background display a uniform prophase arrest (Voelkel-Meiman et al. 2012). We found that in *ndt80Δ*-arrested cells, Pch2-nlsΔ levels were not significantly reduced compared to those of the wild-type Pch2 (Fig. 1f, g), demonstrating that the disappearance of Pch2-nlsΔ at late time points in the *zip1Δ pch2-nlsΔ* double mutant (Fig. 1e) is the consequence, and not the cause, of meiotic cell cycle progression. We also analyzed molecular markers of checkpoint activation influenced by Pch2 function, such as Mec1-dependent Hop1-T318 phosphorylation and Mek1-dependent H3-T11 phosphorylation (Govin et al. 2010; Penedos et al. 2015; Herruzo et al. 2016; Kniewel et al. 2017). The *zip1Δ* mutant showed high levels of these phosphorylation events that were largely abolished in both *zip1Δ pch2Δ* and *zip1Δ pch2-nlsΔ* (Fig. 1e) consistent with suppression of the meiotic block. Thus, the *zip1Δ pch2-nlsΔ* mutant phenocopies the checkpoint defects of *zip1Δ pch2Δ*, indicating that the NLS-like sequence is absolutely required for meiotic checkpoint function.

**Table 1.**
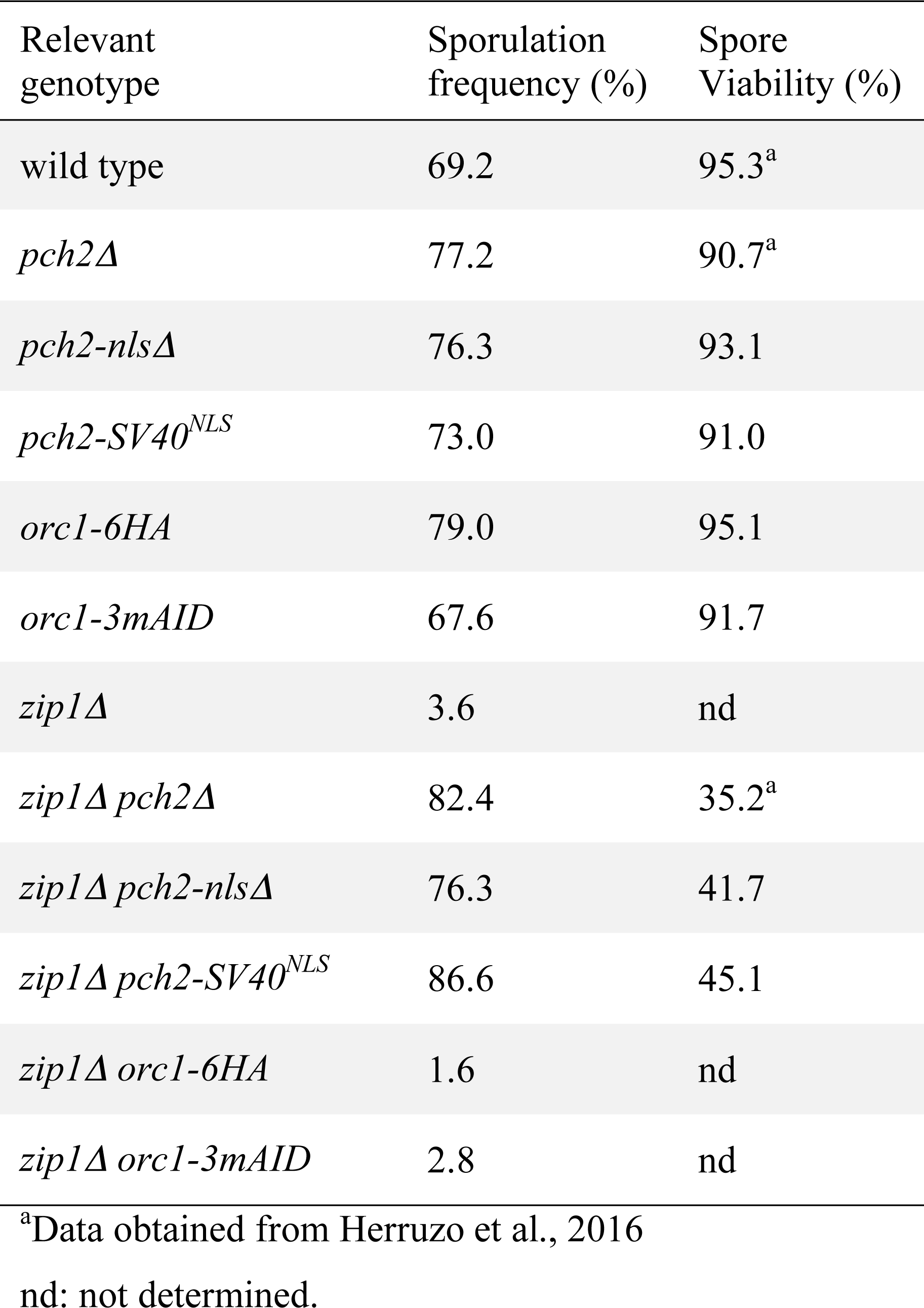
Sporulation and spore viability.

**Fig. 1.**
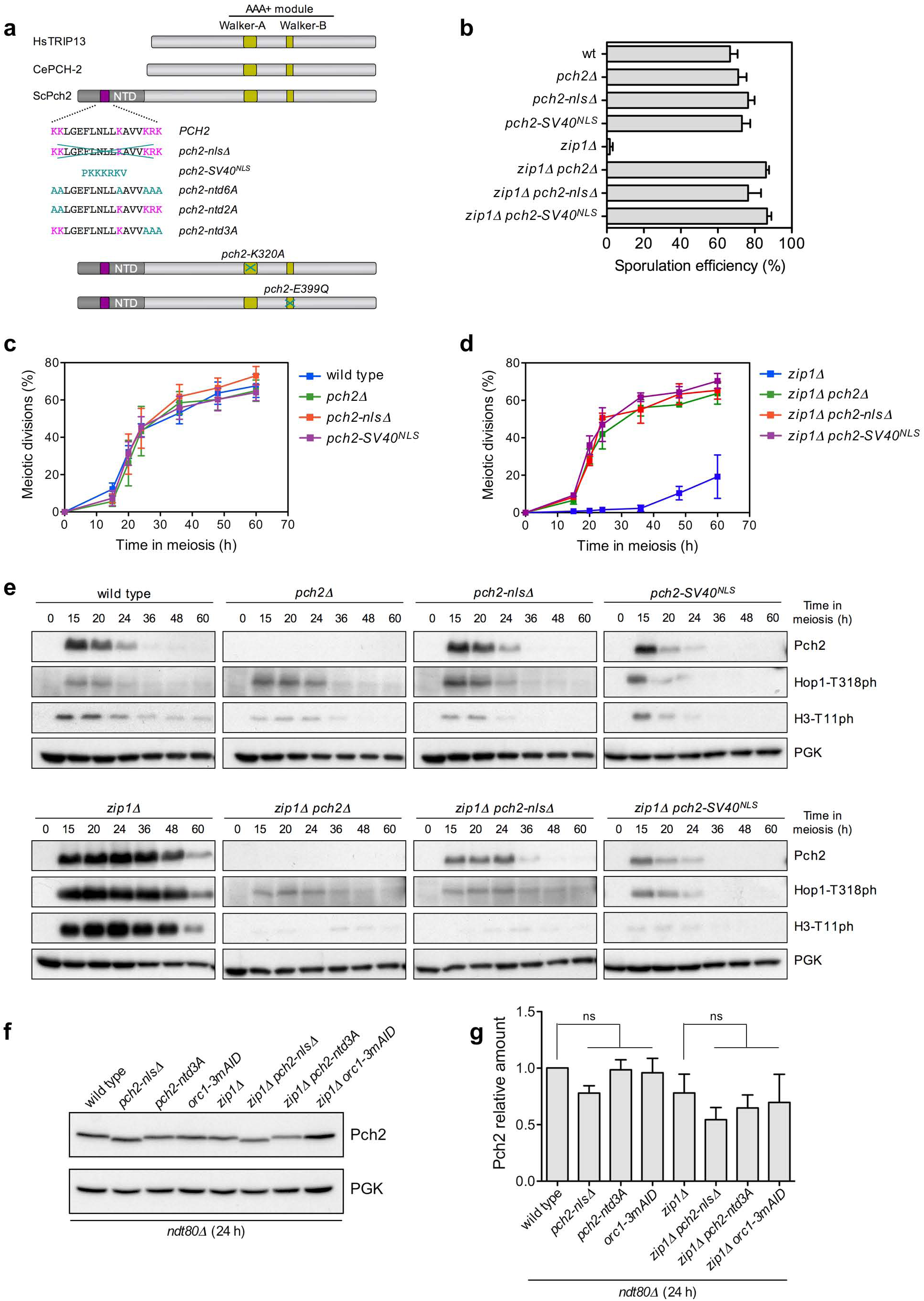
A basic-rich motif in the Pch2 NTD is essential for its checkpoint function. **a** Pch2 relevant motifs and mutants generated. A schematic representation of the *S*. *cerevisiae* Pch2 protein (ScPch2) and the orthologs from *C*. *elegans* (CePCH-2) and human (HsTRIP13) is depicted indicating the characteristic AAA+ ATPase motifs. The sequence of the basic-rich motif (purple) in the extended N-terminal domain of Pch2 (NTD) is shown along with the modifications introduced (light blue) in the different mutants generated in this work (see text). The Walker A and Walker B mutants previously constructed are also shown (Herruzo et al. 2016). **b** Sporulation efficiency, determined by microscopic counting, after 3 days on sporulation plates. Error bars: SD; n=3. **c, d** Time course analysis of meiotic nuclear divisions; the percentage of cells containing two or more nuclei is represented. Error bars: SD; n=6 in (**c**); n=3 in (**d**). **e** Western blot analysis of Pch2 production during meiosis (detected with anti-HA antibodies), Hop1-T318 phosphorylation and Mek1 activation (H3-T11 phosphorylation). PGK was used as a loading control. Strains in (**b, c, d, e**) are: DP1151 (wild type), DP1164 (*pch2*Δ), DP1408 (*pch2-nlsΔ*), DP1455 (*pch2-SV40*^*NLS*^), DP1152 (*zip1Δ*), DP1161 (*zip1Δ pch2Δ*), DP1409 (*zip1Δ pch2-nlsΔ*) and DP1456 (*zip1Δ pch2-SV40*^*NLS*^). **f** Western blot analysis of Pch2 production in *ndt80Δ*-arrested strains of the indicated genotypes. Auxin (500 μ M) was added to *orc1-m3AID* cultures 12 hours after meiotic induction and all cell extracts were prepared at 24 hours. **g** Quantification of Pch2 levels normalized with PGK and relativized to wild type. Errors bars: SD; n=3; ns: not significant. The *ndt80Δ* strains in (**f, g**) are DP1191 (wild type), DP1411 (*pch2-nlsΔ*), DP1569 (*pch2-ntd3A*), DP1451 (*orc1-m3AID*), DP1190 (*zip1Δ*), DP1412 (*zip1Δ pch2-nlsΔ*), DP1570 (*zip1Δ pch2-ntd3A*) and DP1452 (*zip1Δ orc1-3mAID*).

We also examined the localization of the Pch2-nlsΔ protein by immunofluorescence of spread pachytene chromosomes at 24 h after meiotic induction in an *ndt80Δ* background. We used antibodies recognizing the Nsr1 protein as a nucleolar marker. Nuclei with a zygotene/pachytene chromosomal morphology based on DAPI staining of chromatin were scored in all localization analyses. As previously described, in both wild type and *zip1Δ* nuclei, Pch2 displayed a conspicuous accumulation at the rDNA region marked by the presence of Nsr1 in the vicinity (Fig. 2a; Table S1). In contrast, the Pch2-nlsΔ protein was not detected associated to the meiotic rDNA chromatin (Fig. 2a; Table S1) despite the fairly normal levels observed in whole-cell extracts (Fig. 1f, g). Thus, the NLS-like stretch is required for Pch2 nucleolar localization.

**Fig. 2.**
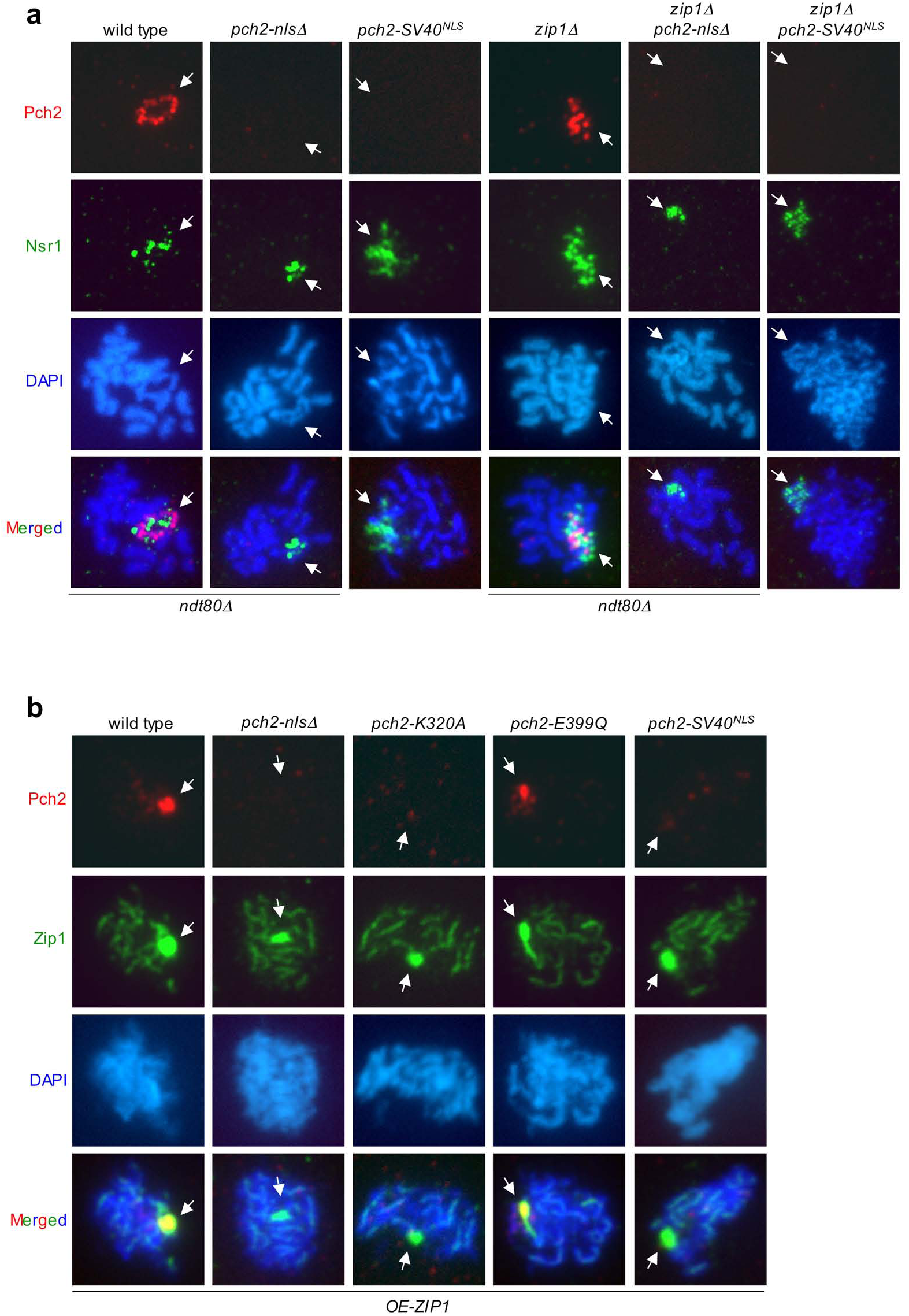
The basic-rich motif in the Pch2 NTD is essential for its nucleolar localization and for its association with SC components. **a** Immunofluorescence of meiotic chromosomes stained with anti-Pch2 antibodies to detect Pch2, Pch2-nlsΔ or Pch2-SV40^NLS^ (red), anti-Nsr1 antibodies (green) and DAPI (blue). Representative nuclei are shown. Arrows point to the rDNA region. Samples were prepared 24 hours after meiotic induction for the *ndt80Δ* strains DP1191 (wild type), DP1411 (*pch2-nls*Δ), DP1190 (*zip1Δ*) and DP1412 (*zip1Δ pch2-nlsΔ*), or at 15 hours for DP1455 (*pch2-SV40*^*NLS*^) and DP1456 (*zip1Δ pch2-SV40*^*NLS*^). **b** Immunofluorescence of meiotic chromosomes stained with anti-HA antibodies to detect Pch2, Pch2-nlsΔ, Pch2-K320A, Pch2-E399Q or Pch2-SV40^NLS^ (red), anti-Zip1 antibodies (green) and DAPI (blue). Representative nuclei are shown. Samples were prepared 15 hours after meiotic induction. Arrows point to the polycomplex. Strains in (**b**) are DP1151 (wild type), DP1408 (*pch2-nls*Δ), DP1163 (*pch2-K320A*), DP1287 (*pch2-E399Q*), and DP1455 (*pch2- SV40*^*NLS*^), all of them transformed with a high-copy plasmid overexpressing *ZIP1* (pSS343).

The faint rDNA-independent SC-associated foci of Pch2 on wild-type prophase chromosomes are difficult to detect in most nuclear spread preparations (at least from BR strains) because their intensity is only slightly above the background level and they are often masked by the intense nucleolar signal (Herruzo et al. 2016). To circumvent this issue, we devised an alternative strategy to monitor the ability of Pch2 (or mutant derivatives) to bind SC components by inducing the formation of polycomplexes. The polycomplex is an extrachromosomal aggregate of SC proteins formed under certain circumstances (i.e., *ZIP1* overexpression or Spo11 deficiency) that mimics the same ultrastructure as the native SC (Dong and Roeder 2000). The formation of this structure provides an excellent and prominent readout for SC assembly allowing us to easily assess Pch2 interaction with SC components (Fig. 2b). Therefore, we examined the presence of different Pch2 versions in the polycomplexes of strains overexpressing *ZIP1*. As expected, the wild-type Pch2 protein extensively colocalized with Zip1 in the polycomplex (San-Segundo and Roeder 1999); in contrast, Pch2-nlsΔ failed to be detected in this structure (Fig. 2b; Table S1). We conclude that the NTD portion deleted in Pch2-nlsΔ is not only necessary for rDNA localization, but also for interaction with SC proteins.

We also analyzed the ability of the checkpoint-deficient ATPase-dead versions of Pch2 previously generated (Pch2-K320A and Pch2-E399Q) (Fig. 1a) to interact with Zip1 in the polycomplex. Pch2-K320A lacks the ATP-binding site in the Walker-A motif; this protein fails to maintain a stable AAA+ hexameric complex and also fails to localize to meiotic chromatin. Pch2-E399Q lacks the ATP-hydrolysis site in the Walker-B motif, but it does localize to the rDNA region despite being catalytically inactive (Chen et al. 2014; Herruzo et al. 2016). Consistent with those observations, we found that Pch2-K320A does not localize to polycomplexes, whereas the Pch2-E399Q version retains the capacity to associate with SC components (Fig. 2b; Table S1). Thus, the ATPase activity of Pch2 is not intrinsically a requisite for its proper localization.

To assess whether the basic-rich 17-amino-acid NTD sequence is actually acting as a true NLS to sustain Pch2 function, we replaced it by a bona-fide NLS from the SV40 virus generating the *pch2-SV40*^*NLS*^ version (Fig. 1a). Albeit with slightly reduced levels, the Pch2-SV40^NLS^ protein displayed the characteristic kinetics of prophase induction and eventual disappearance coincident with meiotic progression similar to Pch2 and Pch2-nlsΔ (Fig. 1e). Like *pch2Δ* and *pch2-nlsΔ*, the *pch2-SV40*^*NLS*^ single mutant sustained normal sporulation and high levels of spore viability (Fig. 1b, Table 1). In addition, similar to *zip1Δ pch2Δ* and *zip1Δ pch2-nlsΔ*, the *zip1Δ*-induced checkpoint-dependent meiotic block was alleviated in the *zip1Δ pch2-SV40*^*NLS*^ double mutant resulting in increased spore death (Fig. 1b, d; Table 1). Consistently, *zip1Δ pch2-SV40*^*NLS*^ displayed impaired Hop1-T318 and H3-T11 phosphorylation as compared to *zip1Δ* (Fig. 1e). Moreover, like Pch2-nlsΔ, the Pch2-SV40^NLS^ version also failed to localize to the rDNA region and to the polycomplex on meiotic chromosome spreads (Fig. 2a, b; Table S1). Thus, these findings are consistent with the possibility that the function of the Pch2 NTD motif is not, at least exclusively, driving Pch2 nuclear or nucleolar targeting/import by a canonical NLS-dependent mechanism.

### The “KRK” basic motif in the Pch2 NTD is required for checkpoint function and localization

In order to pinpoint the residues within the 17-amino acid stretch that are relevant for Pch2’s checkpoint function we constructed several mutants in which the basic residues were changed to alanine in different combinations (Fig. 1a): in *pch2-ntd6A*, all lysines (K) and the arginine (R) were mutated; in *pch2-ntd2A*, the KK at positions 42-43 were mutated and in *pch2-ntd3A*, the KRK at positions 56-58 were mutated. We introduced these mutations into centromeric plasmids containing 3HA-tagged *PCH2* and transformed a checkpoint-deficient *zip1Δ pch2Δ* strain to assess their ability to restore checkpoint function by monitoring sporulation efficiency (Fig. 3a). As controls, the *zip1Δ pch2Δ* strain was also transformed with the empty vector (checkpoint fully inactive) or with the wild-type *PCH2* (checkpoint active). We found that *pch2-ntd6A* and *pch2-ntd3A* completely suppressed the *zip1Δ* sporulation defect, whereas *pch2-ntd2A* only conferred a partial decrease in sporulation efficiency. Note that the wild-type *PCH2* did not fully restore checkpoint arrest in this plasmid-based assay as a consequence of plasmid-loss events. Those *zip1Δ pch2Δ* cells that lose the plasmid (about 25%) become completely checkpoint defective and complete sporulation (Refolio et al. 2011). In any case, these observations suggest that the Pch2-ntd6A and Pch2-ntd3A proteins do not support checkpoint function, but the Pch2-ntd2A version retains partial activity. To confirm this conclusion we analyzed H3-T11 phosphorylation as a reporter for Mek1 activity. Consistent with the meiotic phenotype, H3-T11ph was severely impaired in the strains harboring *pch2-ntd6A* and *pch2-ntd3A* mutations and only partially reduced in *pch2-ntd2A* (Fig. 3b) indicating that the KRK motif at positions 56-58 of the Pch2 NTD is critical for the meiotic recombination checkpoint. Note that the absence of the Pch2-nlsΔ, Pch2-ntd6A and Pch2-ntd3A proteins at the 24-hour time point (Fig. 3b) is the consequence of meiotic progression in these checkpoint-deficient mutants since both Pch2-nlsΔ and Pch2-ntd3A are produced at fairly normal levels in prophase-arrested *ndt80Δ* strains (Fig. 1f, g).

**Fig. 3.**
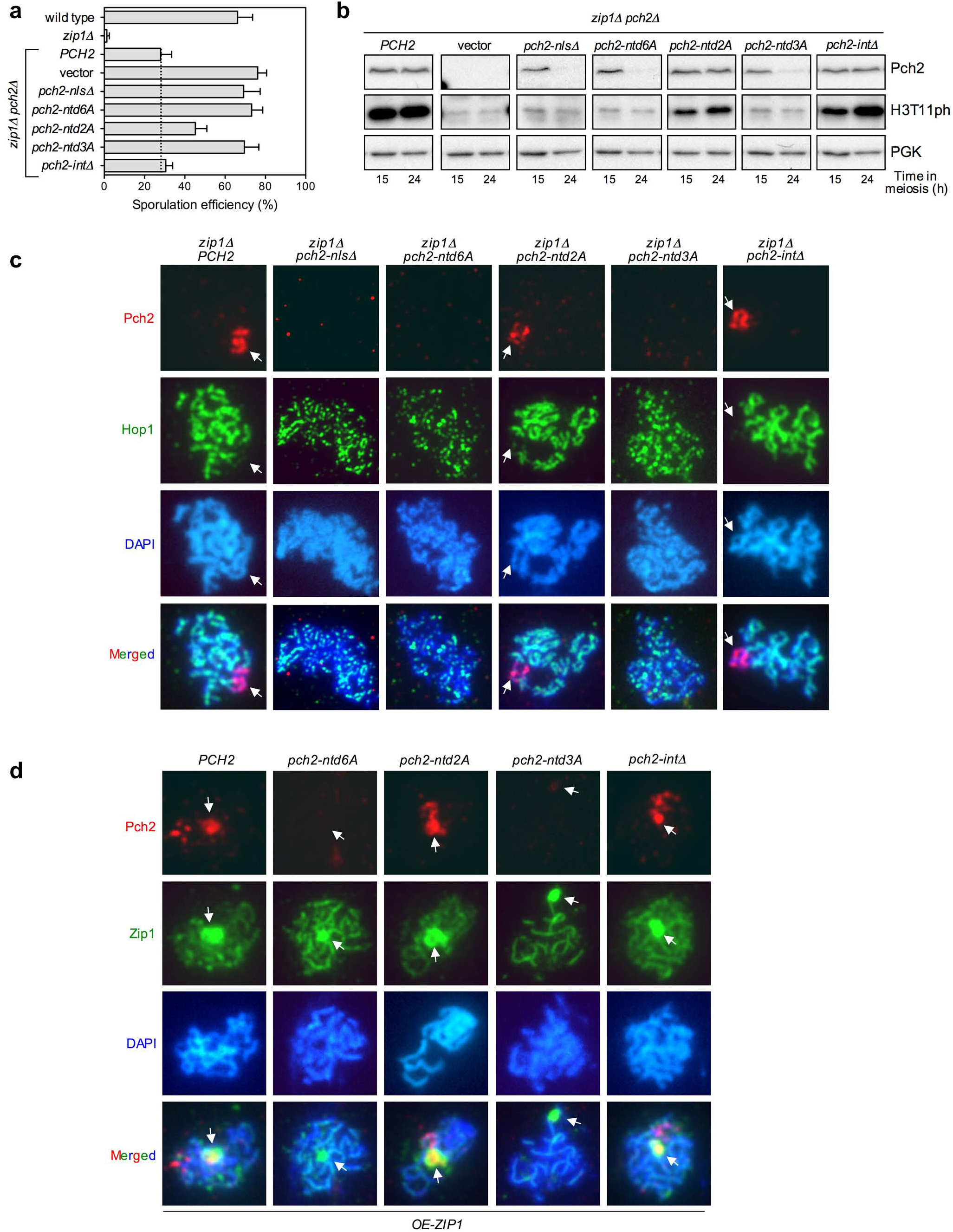
The KRK sequence within the basic motif in the Pch2 NTD is essential for its checkpoint function, nucleolar localization and association with SC components. **a** Sporulation efficiency, determined by microscopic counting, after 3 days on sporulation plates. Error bars: SD; n=3. The dotted line marks the basal level of complementation of *zip1Δ pch2Δ* checkpoint defect with the wild-type *PCH2* plasmid discarding the plasmid-loss effect. **b** Western blot analysis of Pch2 production and Mek1 activation (H3-T11 phosphorylation) at the indicated times after meiotic induction. PGK was used as a loading control. Strains in (**a**, **b**) are: DP421 (wild type), DP422 (*zip1Δ*) and DP1405 (*zip1Δ pch2Δ*). The *zip1Δ pch2Δ* strain was transformed with pSS75 (*PCH2*), pRS314 (vector), pSS338 (*pch2-nlsΔ*), pSS358 (*pch2-ntd6A*), pSS363 (*pch2-ntd2A*), pSS364 (*pch2-ntd3A*) and pSS362 (*pch2-intΔ*). **c** Immunofluorescence of meiotic chromosomes stained with anti-HA antibodies to detect Pch2, Pch2-nlsΔ, Pch2-ntd6A, Pch2-ntd2A, Pch2-ntd3A or Pch2-intΔ (red), anti-Hop1 antibodies (green) and DAPI (blue). Representative nuclei are shown. Samples were prepared 15 hours after meiotic induction. Arrows point to the rDNA region. The DP1405 (*zip1Δ pch2Δ*) strain was transformed with pSS75 (*PCH2*), pSS338 (*pch2-nlsΔ*), pSS358 (*pch2-ntd6A*), pSS363 (*pch2-ntd2A*), pSS364 (*pch2-ntd3A*) and pSS362 (*pch2-intΔ*). **d** Immunofluorescence of meiotic chromosomes stained with anti-HA antibodies to detect Pch2, Pch2-ntd6A, Pch2-ntd2A, Pch2-ntd3A or Pch2-intΔ (red), anti-Zip1 antibodies (green) and DAPI (blue). Representative nuclei are shown. Samples were prepared 15 hours after meiotic induction. Arrows point to the polycomplex. The DP186 (*pch2Δ*) strain, transformed with pSS75 (*PCH2*), pSS358 (*pch2-ntd6A*), pSS363 (*pch2-ntd2A*), pSS364 (*pch2-ntd3A*) and pSS362 (*pch2-intΔ*), was also co-transformed with a high-copy plasmid overexpressing *ZIP1* (pSS343).

We next examined the localization of these Pch2-ntd mutant versions on spread *zip1Δ* meiotic nuclei in combination with Hop1 staining at the 15 h time point when all the proteins are present. Both Pch2-ntd6A and Pch2-ntd3A failed to decorate the rDNA meiotic chromatin. Moreover, like *zip1Δ pch2Δ* (Herruzo et al. 2016), the *zip1Δ pch2-ntd6A and zip1Δ pch2-ntd3A* mutants displayed fragmented and discontinuous Hop1 distribution, in contrast to the linear Hop1 axial configuration characteristic of the *zip1Δ* mutant (Fig. 3c; Table S1). On the other hand, the Pch2-ntd2A protein, which confers partial checkpoint activity, did localize to the rDNA retaining the capacity to exclude Hop1 from the nucleolar region (Fig. 3c; Table S1). We also determined the ability to interact with SC components by analyzing colocalization with Zip1 in polycomplexes. We overexpressed *ZIP1* from a high-copy vector in *pch2Δ* strains co-transformed with centromeric plasmids expressing either wild-type *PCH2* or the different *pch2-ntd* mutant versions. As expected, we observed extensive colocalization of wild-type Pch2 and Zip1 within the polycomplex (Fig. 3d; Table S1). In contrast, Pch2-ntd6A and Pch2-ntd3A did not associate with polycomplexes, whereas Pch2-ntd2A could be detected in this structure (Fig. 3d; Table S1). Thus, both *pch2-ntd6A* and *pch2-ntd3A* mutants, but not *pch2-ntd2A*, appear to be defective in Pch2 localization and meiotic recombination checkpoint function, suggesting that the KRK motif in the context of the Pch2 NTD is crucial for Pch2 action in the response to *zip1Δ*-induced meiotic defects.

### The *PCH2* intron is not relevant for the meiotic checkpoint

The mRNA produced by the *PCH2* gene contains an intron close to the end that undergoes Tgs1-dependent and Mer1-independent splicing (Fig. S2) (Qiu et al. 2011). Although most budding yeast genes do not possess introns, their presence is relatively frequent among meiotic genes; indeed, controlled intron processing is crucial for certain meiotic events (Munding et al. 2010). In order to investigate if the regulated splicing of the *PCH2* mRNA is required for a proper meiotic checkpoint response, we constructed a centromeric plasmid carrying a *PCH2* allele lacking the intron sequence (*pch2-intΔ*) (Fig. S2) and assessed its ability to restore checkpoint function when transformed into a *zip1Δ pch2Δ* mutant. The Pch2 protein was produced from the *pch2-intΔ* allele with similar dynamics as the protein produced from the wild-type *PCH2* gene (Fig. 3b). Introduction of the *pch2-intΔ* allele decreased sporulation efficiency of *zip1Δ pch2Δ* to the same levels as the wild-type *PCH2* did (Fig. 3a), and it sustained high levels of H3-T11 phosphorylation (Fig. 3b). Moreover, like the protein produced from the wild-type *PCH2* gene, the Pch2 protein generated from the *pch2-intΔ* allele localized to the rDNA excluding Hop1 from this region (Fig. 3c; Table S1) and also was capable of interacting with Zip1 in the polycomplex (Fig. 3d; Table S1). Therefore, although we cannot rule out a subtle effect in other meiotic events controlled by Pch2, we conclude that the *PCH2* intron is dispensable for the *zip1Δ*-induced meiotic recombination checkpoint and for Pch2 chromosomal localization.

### Analysis of Pch2 localization in whole meiotic cells

Using chromosome spreading, we have shown above that deletion or mutation of the NLS-like motif in the Pch2 NTD prevents its rDNA localization and association with SC proteins also leading to defective checkpoint function. Insertion of a bona-fide NLS from SV40 neither restores Pch2 chromosome binding nor function. Therefore, to further investigate the contribution of this basic NTD motif to govern Pch2 location, we explored Pch2 subcellular localization in whole meiotic cells. For this purpose, we initially generated diploid strains (*GFP-PCH2*) expressing a version of the *PCH2* gene containing the sequence of the green fluorescent protein (GFP) inserted at the second codon in its own genomic locus. A flexible linker encoding five Gly-Ala repeats was also introduced between the *GFP* and *PCH2* sequences (Fig. S3a). Several lines of evidence demonstrated that the GFP-Pch2 protein is functional. First, *zip1Δ GFP-PCH2* strains displayed a tight sporulation block similar to that of *zip1Δ* (Fig. S3b). Second, like the *zip1Δ* mutant, *zip1Δ GFP-PCH2* showed a marked meiotic delay in meiotic time courses (Fig. S3c), and sustained Hop1-T318 and H3-T11 phosphorylation (Figure S3d). Third, Hop1 was excluded from the rDNA region in *zip1Δ GFP-PCH2* meiotic chromosomes (Figure S3e). Nevertheless, despite being functional, western blot analysis revealed that GFP-Pch2 was produced in meiotic cells at about 10-fold reduced levels compared to the wild-type Pch2 protein and was barely detectable (Fig. 4a; second lane; Fig. S3d). These observations indicate that extremely low levels of Pch2 are sufficient to establish the meiotic checkpoint response, but prevent the use of the endogenous *GFP-PCH2* fusion for precise and sensitive localization studies. Thus, we placed the *GFP-PCH2* construct (as well as *GFP-pch2-nlsΔ* and *GFP-pch2-SV40*^*NLS*^) under control of the meiosis-specific *HOP1* promoter in centromeric plasmids (Fig 4b). In this situation, GFP-Pch2 and GFP-Pch2-nlsΔ were produced at roughly similar levels as the untagged Pch2 protein, and GFP-Pch2-SV40^NLS^ at only slightly reduced levels (Fig. 4a). Moreover, this plasmid-borne version of *GFP-PCH2* was capable of restoring sporulation arrest to large extent when transformed into *zip1Δ pch2Δ* (Fig. 4c) (sporulation was not completely blocked due to plasmid-loss events; see previous sections for explanation). In contrast, GFP-Pch2-nlsΔ and GFP-Pch2-SV40^NLS^ did not confer checkpoint functionality (Fig. 4c), consistent with the results shown above (Fig. 1b, d). Consequently, we used these constructs to examine Pch2 distribution in whole meiotic prophase cells. These plasmids were transformed into *zip1Δ* strains, also harboring *HOP1-mCherry* in heterozygosis as a marker for meiotic prophase chromosomes, and were analyzed by fluorescence microscopy.

**Fig. 4.**
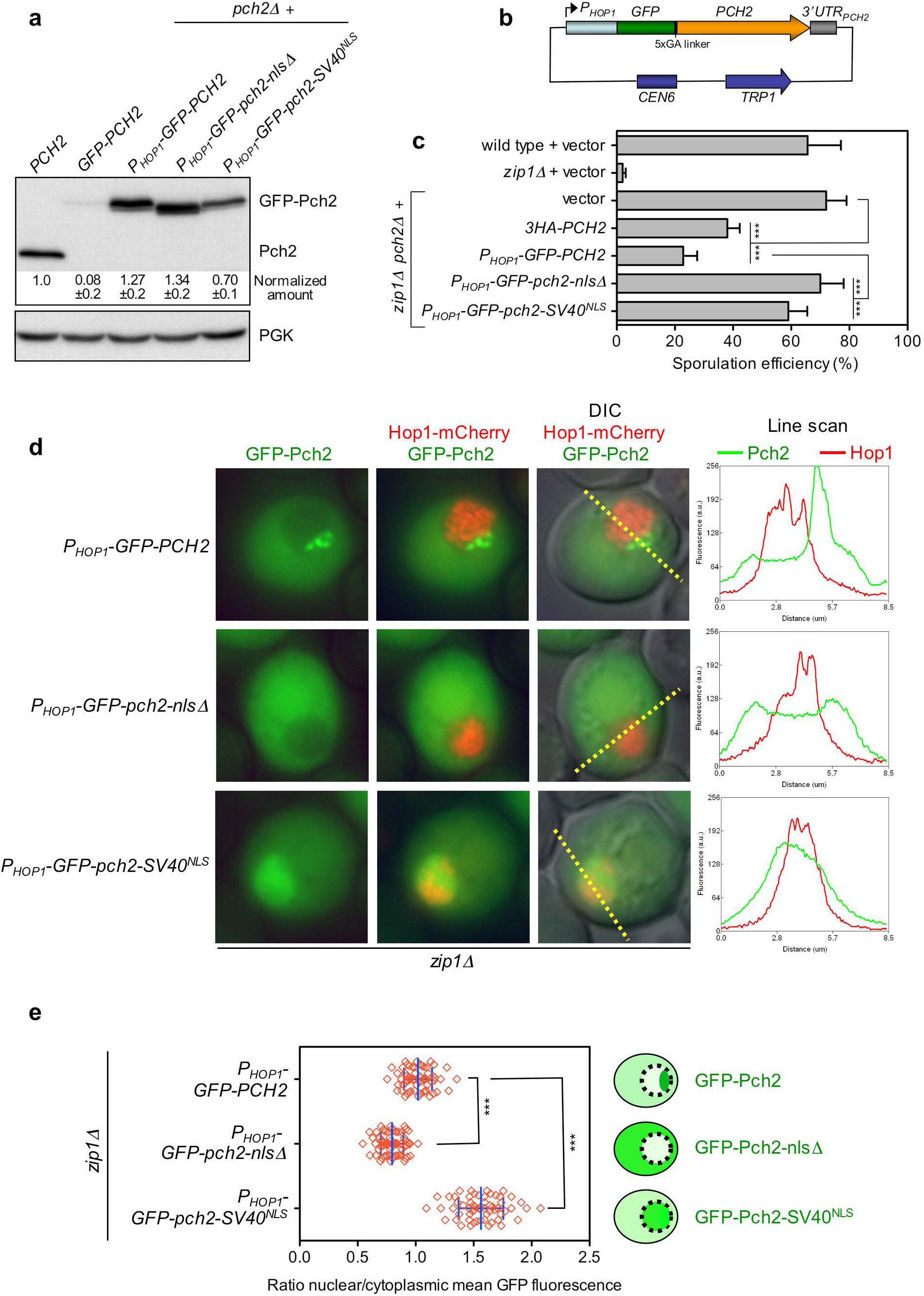
The basic NLS-like motif in Pch2 NTD orchestrates its proper subcellular distribution. **a** Production of untagged Pch2 and different versions of GFP-Pch2 was analyzed by western blot 15 hours after meiotic induction. Protein levels were normalized with PGK and relativized to untagged wild-type Pch2; n=4. Strains are BR2495 (*PCH2*), DP1508 (*GFP-PCH2*) and DP186 (*pch2Δ*). DP186 was transformed with pSS393 (*P*_*HOP1*_*-GFP-PCH2*), pSS396 (*P*_*HOP1*_*-GFP-pch2-nlsΔ*) and pSS397 (*P*_*HOP1*_*-GFP-pch2-SV40*^*NLS*^). **b** Schematic representation of the *P*_*HOP1*_*-GFP-PCH2* construct in the pSS393 plasmid. The pSS396 and pSS397 plasmids (not depicted) are similar, but express *GFP-pch2-nlsΔ* and *GFP-pch2-SV40*^*NLS*^, respectively. **c** Sporulation efficiency, determined by microscopic counting, after 3 days on sporulation plates. Error bars: SD; n=6. Strains are DP421 (wild type), DP422 (*zip1Δ*) and DP1405 (*zip1Δ pch2Δ*), transformed with pRS314 (vector), pSS75 (*3HA-PCH2*), pSS393 (*P*_*HOP1*_*-GFP-PCH2*), pSS396 (*P*_*HOP1*_*-GFP-pch2-nlsΔ*) or pSS397 (*P*_*HOP1*_*-GFP-pch2-SV40*^*NLS*^), as indicated. **d** Fluorescence microscopy analysis of GFP-Pch2 (green) and Hop1-mCherry (red) distribution in whole meiotic cells 15 hours after meiotic induction. The overlay with differential interference contrast (DIC) images is also displayed to show the cell morphology. The plots represent the GFP and mCherry fluorescent signals (green and red, respectively) along the depicted yellow lines from left to right. Representative cells are shown. Additional cells and line-scan plots are presented in Fig. S4. **e** Quantification of the ratio between the nuclear (including the nucleolar) and cytoplasmic GFP fluorescent signal. The cartoon illustrates the subcellular localization of the different Pch2 versions. The strains in (**d**, **e**) are DP1500 (*zip1Δ*) transformed with pSS393 (*P*_*HOP1*_*-GFP-PCH2*), pSS396 (*P*_*HOP1*_*-GFP-pch2-nlsΔ*) or pSS397 (*P*_*HOP1*_*-GFP-pch2-SV40*^*NLS*^); 62, 72 and 59 cells, respectively, were scored.

We found that the wild-type GFP-Pch2 localized mainly to a discrete reduced area in one side of the nucleus (Fig. 4d; Fig. S4). According with the prominent localization pattern of Pch2 on chromosome spreads (Fig. 2a) (San-Segundo and Roeder 1999; Herruzo et al. 2016) and with the fact that this conspicuous GFP-Pch2 structure did not overlap with Hop1-mCherry (Fig. 4d; Fig. S4) we conclude that it likely corresponds to the nucleolus. In addition, GFP-Pch2 also displayed a diffuse homogenous cytoplasmic signal (Fig. 4d; Fig. S4). In contrast, GFP-Pch2-nlsΔ was largely excluded from the nucleus and found mostly in the cytoplasm (Fig. 4d; Fig. S4), as demonstrated by the reduced nuclear/cytoplasm fluorescence ratio of *GFP-pch2-nlsΔ* cells compared to that of wild-type *GFP-PCH2* (Fig 4e). On the other hand, GFP-Pch2-SV40^NLS^ was more concentrated inside the nucleus (Fig. 4e) displaying a diffuse nucleoplasmic signal, but did not show nucleolar accumulation (Fig. 4d; Fig. S4). The use of the LineScan tool of MetaMorph software to trace fluorescent signals confirmed the differential distribution of GFP-Pch2, GFP-Pch2-nlsΔ and GFP-Pch2-SV40^NLS^ across nucleolar, nuclear and cytoplasmic compartments (Fig. 4d; Fig. S4).

These results indicate that the basic-rich motif in the NTD of Pch2 is required for its nuclear/nucleolar accumulation, but it is not simply acting as a canonical NLS sequence. The substitution of this motif for the SV40 NLS is capable of bringing Pch2 back to the nucleus, but it does not restore its normal distribution or its checkpoint function. We conclude that Pch2’s NTD basic motif drives Pch2 subcellular localization and function by additional mechanisms besides the mere control of nuclear import.

### Orc1 and Pch2 colocalize in the nucleolar region

The results presented above allowed us to identify a short motif in the Pch2 NTD important for its function and localization. We next sought for possible Pch2-interacting factors that could orchestrate Pch2 chromosomal distribution to support its checkpoint role. It has been described that Orc1 interacts with Pch2 promoting its nucleolar targeting to exert a repressive effect on meiotic DSB formation in the rDNA region (Vader et al. 2011). However, the possible implication of Orc1 in the meiotic recombination checkpoint remains to be tested. We first analyzed the localization of Pch2 and Orc1 on spread preparations of meiotic chromosomes. In order to detect Orc1 we constructed a C-terminal 6HA-tagged version of the protein. The *ORC1-6HA* strain (also carrying *3MYC-PCH2*) sporulated to normal levels and displayed high levels of spore viability (Fig. S5a; Table 1). Moreover, the *zip1Δ ORC1-6HA 3MYC-PCH2* diploid showed a strong sporulation block (Fig. S5a) indicating that Orc1 tagging does not disturb the meiotic checkpoint response. Immunofluorescence analysis of spread nuclei revealed that Pch2 and Orc1 at least partially colocalize in the rDNA region (Fig. 5a; arrows). Nevertheless, we note that Pch2 was somewhat mislocalized from the nucleolar area in the strain harboring Orc1-6HA, displaying an additional chromosomal punctate pattern that was not observed in Orc1-untagged nuclei (Fig. 5a; arrowheads; Table S1). These and other observations with additional Orc1-tagging attempts (data not shown; see below) indicate that this essential protein appears to be extremely sensitive to structural alterations produced by the fusion to ectopic epitopes. Although the essential replicative function of Orc1-6HA likely remains intact (growth and sporulation are normal in the tagged strains), other functions, such as Pch2 localization, appear to be slightly affected without compromising checkpoint functionality. To corroborate that the nuclear location where Orc1 colocalizes with Pch2 corresponds to the nucleolus, we took advantage of the fact that acetylation of histone H4 at lysine 16 (H4K16ac) is absent from the rDNA region (Cavero et al., 2016). As shown in Fig. 5b, Orc1 accumulation occurred in a region completely devoid of H4K16ac confirming that it coincides with the nucleolus.

**Fig. 5.**
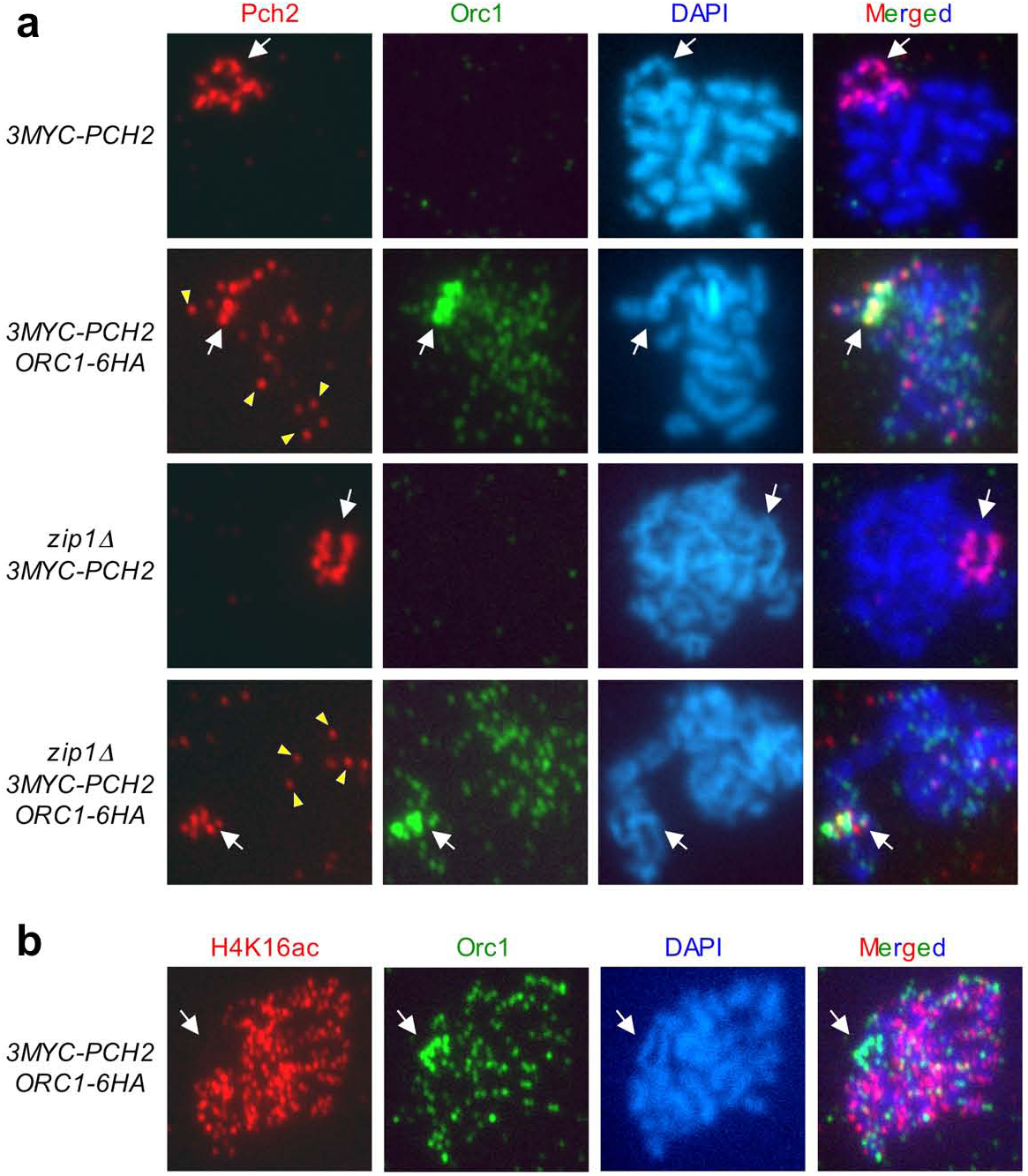
Colocalization of Pch2 and Orc1 in the rDNA region. **a** Immunofluorescence of meiotic chromosomes stained with anti-Pch2 antibodies to detect Pch2 (red), anti-HA antibodies to detect Orc1 (green) and DAPI (blue). Representative nuclei are shown. Samples were prepared 15 hours after meiotic induction. Arrows point to the rDNA region. Arrowheads point to some of the extranucleolar Pch2 dots observed in *ORC1-6HA* nuclei. Strains are DP1243 (*3MYC-PCH2*), DP1426 (*3MYC-PCH2 ORC1-6HA*), DP1244 (*zip1Δ 3MYC-PCH2*) and DP1427 (*zip1Δ 3MYC-PCH2 ORC1-6HA*). **b** Immunofluorescence of meiotic chromosomes stained with anti-H4K16ac (red), anti-HA antibodies to detect Orc1 (green) and DAPI (blue). A representative nucleus is shown. Samples were prepared 15 hours after meiotic induction. Arrows point to the rDNA region. The strain is DP1426 (*3MYC-PCH2 ORC1-6HA*).

### Nucleolar localization of Pch2, but not polycomplex association, is impaired in Orc1-depleted cells

Since *ORC1* is an essential gene, in order to investigate in detail the requirement for Orc1 to target Pch2 to the rDNA and/or to the SC and the implication in the checkpoint, we aimed to generate conditional *orc1* alleles using the auxin-inducible degron (AID) system (Nishimura and Kanemaki 2014). We initially fused the C terminus of Orc1 either to the original full degron tag (AID), a shorter version (mAID), or three tandem copies of it (3mAID), in haploid strains expressing plant *TIR1* from the *ADH1* promoter and assessed the ability to grow on plates containing auxin (Fig. S5b, c). Only the *orc1-3mAID* mutant showed auxin-dependent growth inhibition (Fig. S5c); therefore, we selected this *orc1-3mAID* construct to generate diploid strains harboring the *TIR1* gene under control of the meiosis-specific *HOP1* promoter to use this system for depleting Orc1 in meiotic cultures. The *orc1-3mAID* mutant sustained normal levels of sporulation and spore viability and, like *zip1Δ*, the *zip1Δ orc1-3mAID* double mutant showed a tight sporulation block (Table 1; Fig. S5d). To explore the consequences of Orc1-3mAID depletion in meiotic time courses, we added auxin (or ethanol, as the solvent control) 12 hours after meiotic induction, coinciding with prophase initiation in the BR strain background. We found that Orc1-3mAID was indeed efficiently degraded upon auxin treatment, but Pch2 global levels were not altered when Orc1 was depleted (Fig. 6a and Fig. 1f, g). Analysis of chromosome spreads revealed that, consistent with a previous report using an *orc1-161* thermosensitive allele (Vader et al. 2011), Pch2 was not detected in the nucleolar region of *orc1-3mAID* nuclei either in the presence or absence of added auxin (Fig. 6b; Table S1). This result confirms that Orc1 is required for nucleolar targeting of Pch2 and reveals that C-terminal tagging of Orc1 with the 3mAID degron impairs this particular function without altering other essential roles of Orc1. Thus, Pch2 localization in the rDNA is exquisitely sensitive to Orc1 integrity. In any case, although *orc1-3mAID per se* prevents Pch2 normal distribution, we performed all the ensuing experiments involving this allele with auxin addition to promote Orc1-3mAID degradation (Fig. 6a) and using the untagged version as control thus avoiding uncertainties in the conclusions.

**Fig. 6.**
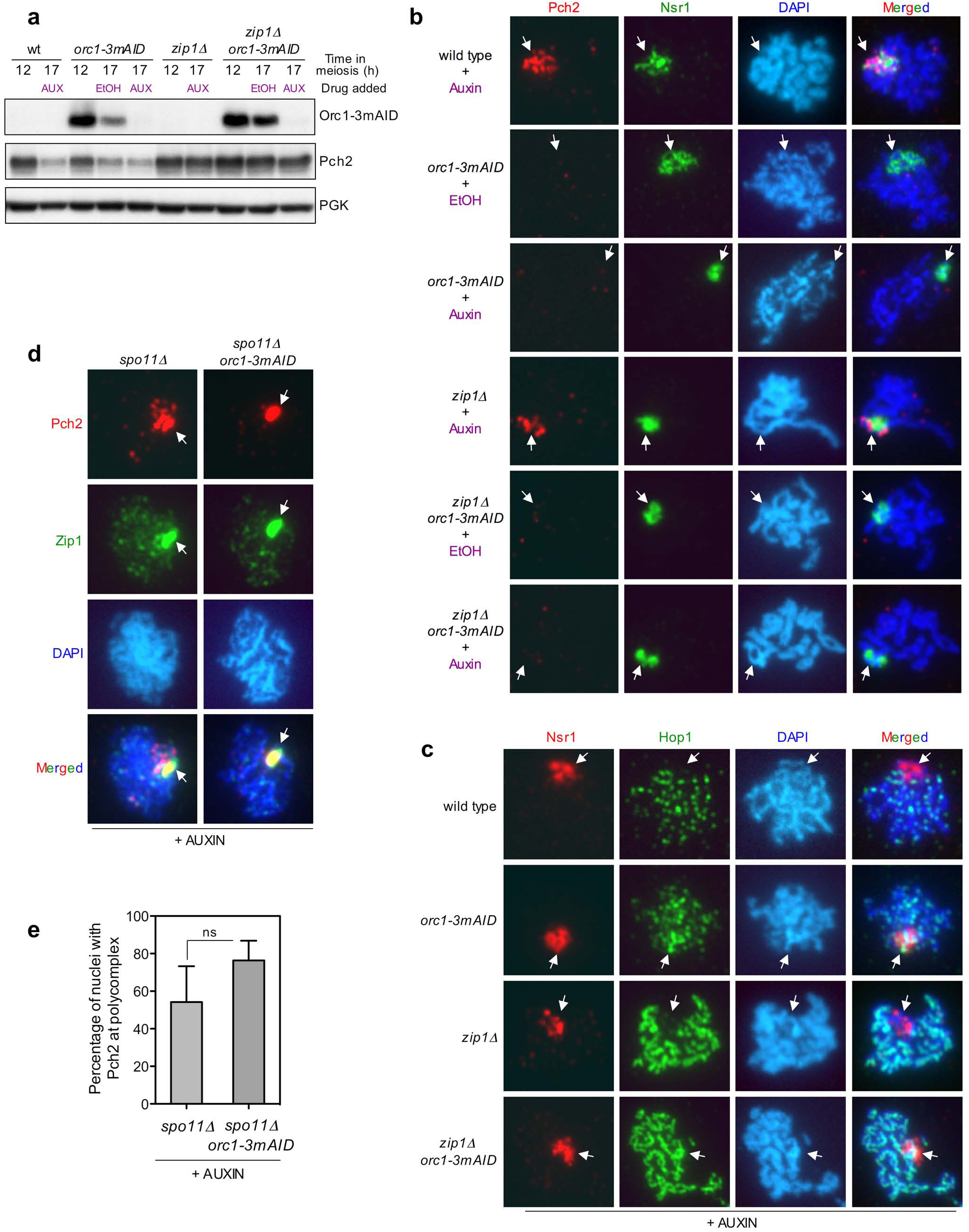
Nucleolar localization of Pch2, but not interaction with SC components, depends on Orc1. **a** Western blot analysis of Pch2 production (detected with anti-HA antibodies) and Orc1-m3AID (detected with anti-mAID antibodies) at the indicated times in meiosis. Auxin (500 μ M) or ethanol (as control) was added 12 hours after meiotic induction. PGK was used as a loading control. **b** Immunofluorescence of meiotic chromosomes stained with anti-Pch2 antibodies to detect Pch2 (red), anti-Nsr1 antibodies (green) and DAPI (blue). Representative nuclei are shown. Auxin (500 μ M) or ethanol (as control) was added 12 hours after meiotic induction and samples were prepared at 17 hours. Arrows point to the rDNA region. **c** Orc1 prevents Hop1 localization to the rDNA. Immunofluorescence of meiotic chromosomes stained with anti-Nsr1 antibodies (red), anti-Hop1 antibodies (green) and DAPI (blue). Representative nuclei are shown. Auxin (500 μ M) was added 12 hours after meiotic induction and samples were prepared at 17 hours. Arrows point to the rDNA region. Strains in (**a**, **b, c**) are DP1151 (wild type), DP1437 (*orc1-3mAID*), DP1152 (*zip1Δ*) and DP1438 (*zip1Δ orc1-3mAID*). **d** Orc1 is dispensable for Pch2 association with the polycomplex. Immunofluorescence of meiotic chromosomes stained with anti-HA antibodies to detect Pch2 (red), anti-Zip1 antibodies (green) and DAPI (blue). Representative nuclei are shown. Auxin (500 μM) was added 12 hours after meiotic induction and samples were prepared at 17 hours. Arrows point to the polycomplex. **e** Quantification of the nuclei displaying Pch2 in the Zip1-containing polycomplex. Error bars, SD; n=3; ns, not significant. Strains in (**d**, **e**) are DP1425 (*spo11Δ*) and DP1444 (*spo11Δ orc1-3mAID*).

Since Pch2 prevents Hop1 binding to the rDNA, we examined the impact of Orc1 depletion on Hop1 localization in both wild-type and *zip1Δ* cells. Consistent with the absence of nucleolar Pch2 in the *orc1-3mAID* mutant (Fig. 6b), Hop1 decorated the rDNA region distinguished by the Nsr1 nucleolar marker (Fig. 6c; Table S1). These observations suggest that Pch2/Orc1-dependent exclusion of Hop1 from the rDNA likely underlies the meiotic DSB repressive effect in this region. We next determined the ability of Pch2 to bind to SC components in the absence of Orc1 by analyzing the colocalization with Zip1 in the polycomplex. In an initial attempt to induce the formation of polycomplexes by overexpressing *ZIP1* from a high-copy plasmid (see above) we found that the *orc1-3mAID* allele precludes *ZIP1* overexpression (Fig. S6), likely as a consequence of a fully functional Orc1 requirement for plasmid maintenance (Fox et al. 1995). Therefore, we took advantage of the recombination and synapsis-defective *spo11Δ* mutant as an alternative tool to promote polycomplex formation (Cheng et al. 2006). In contrast to the *pch2-ntd* mutants characterized above, we observed that Pch2 does colocalize with Zip1 in the polycomplexes formed in the *orc1-3mAID* mutant treated with auxin (Fig. 6d; Table S1). The presence of Pch2 in the *spo11Δ*-induced polycomplexes tended to be even more frequent in *orc1-3mAID* compared to the wild type, although the statistical difference was not significant (Fig. 6e). In sum, these observations indicate that, unlike the rDNA, Pch2 interaction with SC components does not involve Orc1. Thus, the *orc1-3mAID* allele provides a unique scenario to determine whether Pch2 nucleolar localization is specifically required for the meiotic recombination checkpoint response.

### The *zip1Δ* -triggered meiotic recombination checkpoint is active in the absence of Orc1

We analyzed the impact of auxin-induced Orc1-3mAID depletion on the meiotic recombination checkpoint during meiotic time courses (Fig. 7). The *orc1-3mAID* single mutant displayed normal kinetics of nuclear divisions and, more important, like *zip1Δ*, the *zip1Δ orc1-3mAID* double mutant showed a strong delay in meiotic progression (Fig. 7a) suggesting that the checkpoint remains functional in the absence of Orc1. Moreover, the meiotic block of *zip1Δ orc1-3mAID* was alleviated by deletion of *PCH2* (Fig. 7b) indicating that it stems from activation of the meiotic recombination checkpoint. To validate this interpretation, we analyzed Hop1-T318 and H3-T11 phosphorylation as markers of checkpoint activation. In *zip1Δ orc1-3mAID* cells treated with auxin to induce Orc1-m3AID degradation (Fig. 7c), we found high levels of these checkpoint phospho-targets (Fig. 7d), confirming that Orc1 is dispensable for activation and maintenance of the meiotic checkpoint. In addition, to avoid the influence of the different kinetics of meiotic progression of the strains examined, we also quantified the ratio of phospho-Hop1-T318/total Hop1 as a selective indicator of Pch2 checkpoint function in *ndt80Δ*-arrested cells harboring various *pch2* and *orc1* mutations (Herruzo et al. 2016) (Fig. 7e, 7f). We found that, like in *zip1Δ pch2Δ*, the relative levels of Hop1-T318 phosphorylation were drastically reduced in *zip1Δ pch2-nlsΔ* and *zip1Δ pch2-ntd3A*, accounting for the defective Mek1 activation and according with the results presented above; in contrast, Hop1-T318 levels were not significantly altered in *zip1Δ orc1-3mAID*, confirming that the checkpoint was not abrogated. In sum, we can conclude that the Orc1-dependent conspicuous presence of Pch2 in the rDNA region is not required for its function in triggering the meiotic recombination checkpoint, namely sustaining Hop1-T318 phosphorylation.

**Fig. 7.**
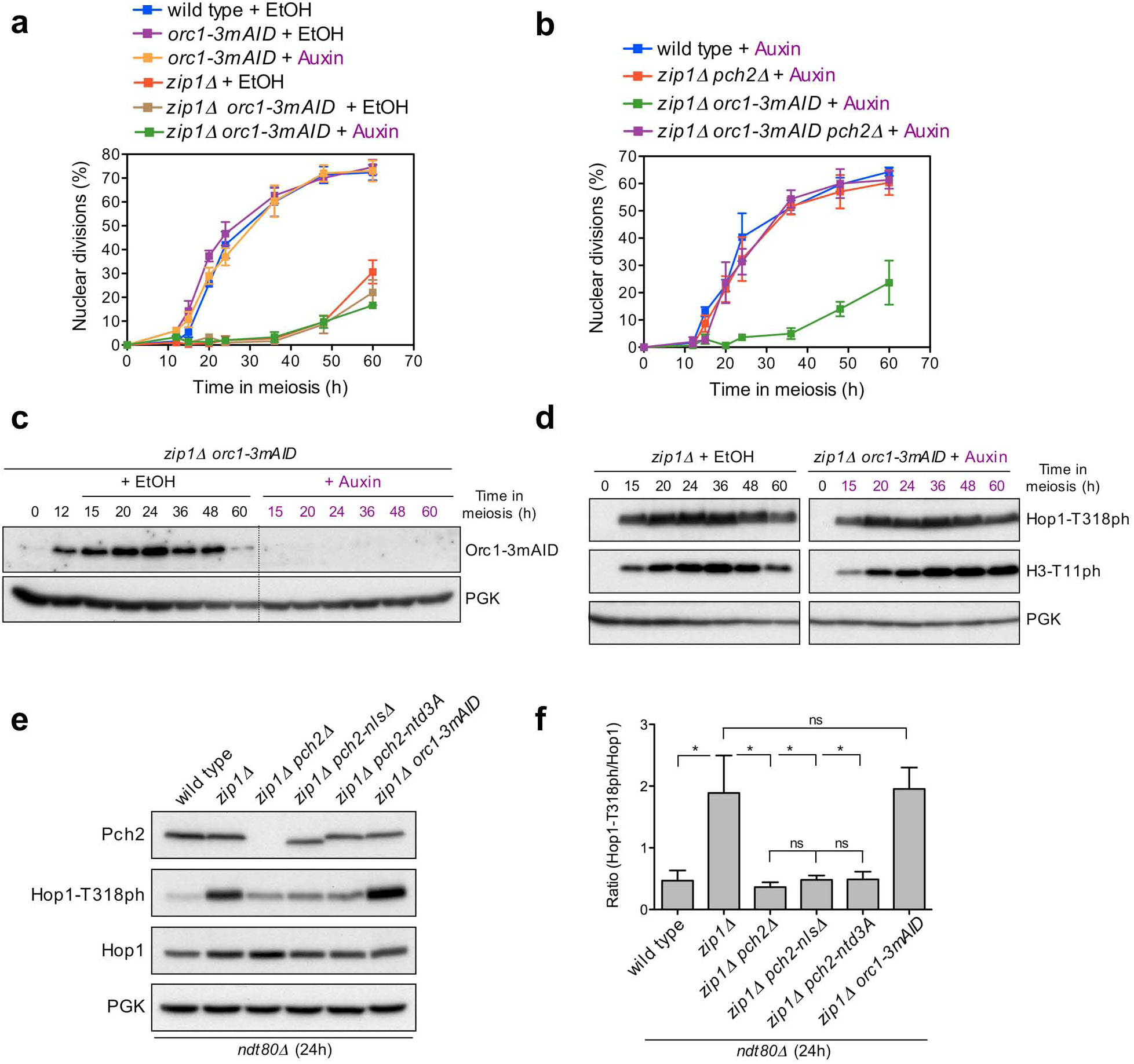
Orc1 is not required for activation of the meiotic recombination checkpoint. **a, b** Time course analysis of meiotic nuclear divisions; the percentage of cells containing two or more nuclei is represented. Error bars: SD; n=3. **c** Western blot analysis of Orc1-m3AID production detected with anti-mAID antibodies. **d** Western blot analysis of Hop1-T318 phosphorylation and Mek1 activation (H3-T11 phosphorylation). In (**a, b, c, d**), auxin (500 μ M) or ethanol (as control) was added 12 hours after meiotic induction. PGK was used as a loading control. Strains in (**a, b, c, d**) are DP1151 (wild type), DP1152 (*zip1Δ*), DP1437 (*orc1-3mAID*), DP1438 (*zip1Δ orc1-3mAID*), DP1161 (*zip1Δ pch2Δ*) and DP1586 (*zip1Δ orc1-3mAID pch2Δ*). **e** Western blot analysis of Pch2 and Hop1 production, and Hop1-T318 phosphorylation in *ndt80Δ*-arrested strains of the indicated genotypes. Auxin (500 μ M) was added to the *zip1Δ orc1-m3AID* culture 12 hours after meiotic induction and all cell extracts were prepared at 24 hours. **f** Quantification of relative Hop1-T318 phosphorylation analyzed as in (**e**). The ratio of phospho-Hop1-T318 versus total Hop1 is represented. Errors bars: SD; n=3. Asterisk: p<0.05; ns: not significant. The *ndt80Δ* strains in (**e**) and (**f**) are DP1191 (wild type), DP1190 (*zip1Δ*), DP881 (*zip1Δ pch2Δ*), DP1412 (*zip1Δ pch2-nlsΔ*), DP1570 (*zip1Δ pch2-ntd3A*) and DP1452 (*zip1Δ orc1-3mAID*).

## DISCUSSION

Previous work has spotted Pch2 as a crucial player in the meiotic recombination checkpoint triggered by the defects provoked by the absence of SC components, such as Zip1 and others (Sym et al. 1993; San-Segundo and Roeder 1999; Wu and Burgess 2006; Herruzo et al. 2016). Pch2 is critically required to sustain high levels of Mec1-dependent Hop1-T318 phosphorylation in order to relay the checkpoint signal to the downstream Mek1 effector kinase. Strikingly, cytological studies reveal that under checkpoint-inducing conditions, such as in the *zip1Δ* mutant lacking the central region of the SC, Pch2 is only detected in the rDNA region raising the possibility that Pch2 exerts its checkpoint function from this particular location. Consistent with this notion, mutations in certain chromatin modifiers that provoke Pch2 mislocalization from the rDNA impair the meiotic checkpoint (San-Segundo and Roeder 2000; Ontoso et al. 2013; Cavero et al. 2016) and also recombination control in perturbed meiosis (Börner et al. 2008). The rDNA array in chromosome XII of budding yeast possesses a unique heterochromatin-like structure that represses recombination (Gottlieb and Esposito 1989). In the case of meiosis, SC formation does not occur in the rDNA and Hop1 binding is prevented (Smith and Roeder 1997). Thus, previous to this study, a puzzling question in the field was how the nucleolar Pch2 could control the phosphorylation status of the axial component Hop1 that is particularly absent in the rDNA region. A paradigm of a crucial cell-cycle regulator governed by the nucleolus is the Cdc14 phosphatase; controlled release of Cdc14 from the nucleolar RENT complex impinges on various processes such as mitotic exit (Stegmeier and Amon 2004), DNA repair (Villoria et al. 2017) and meiotic chromosome segregation (Fox et al. 2017). By analogy with this mechanism, it was possible to speculate that Pch2 may orchestrate the timely nucleolar sequestration and/or release of a critical factor involved in Hop1 phosphorylation.

In order to directly assess the requirement for the nucleolar Pch2 in the meiotic recombination checkpoint, we have identified and characterized *cis* and *trans* localization and functional determinants of Pch2. Table 2 summarizes our findings. The extended NTD of Pch2 was an opportune element to dissect, because it is not conserved in the Pch2 orthologs of other species where the nucleolar localization has not been reported. We have pinpointed a short stretch in Pch2’s NTD containing a KRK basic motif that is essential for its checkpoint function. Nevertheless, this motif is not specific for Pch2 nucleolar targeting; it is also required for interaction with SC components raising the possibility that this basic amino acid stretch may direct global binding of Pch2 to meiotic chromosomes. Alternatively, it was also possible that the only function of this motif is to drive Pch2 nuclear import. The functional analysis of a Pch2-SV40^NLS^ version combined with cytological studies, both in spread chromosomes and whole meiotic cells, has allowed us to address this question. Consistent with the observation that the Pch2-nlsΔ protein does not associate with chromatin on nuclear spreads, it is largely excluded from the nucleus displaying a prominent cytoplasmic localization. Substitution of the NLS-like region by a well-defined NLS from the SV40 virus is capable of promoting the transport of Pch2 to the nucleus but, remarkably, it shows a diffuse nucleoplasmic distribution and, unlike the wild-type Pch2, does not accumulate in the nucleolus. Thus, the fact that Pch2-SV40^NLS^ neither restores Pch2 function nor chromosomal association, despite being present inside the nucleus, implies that the NLS-like region in Pch2’s NTD is not solely acting in Pch2 nuclear transport. This basic patch, which is located in a predicted alpha-helical structure (Fig. S7), may be involved in the interaction of Pch2 with additional factors required for its proper localization and/or function. The N-terminal domain of the Xrs2 protein, a component of the MRX complex, interacts with Pch2 to modulate the Tel1-dependent checkpoint response to unresected meiotic DSBs (Ho and Burgess 2011). Since Tel1 is not required for the *zip1Δ*-induced meiotic block, it is possible to speculate that Xrs2 acts together with Pch2 in the checkpoint response induced by the absence of Zip1 without involving the MRX complex. In fact, MRX-independent functions of Xrs2 have been described in the vegetative DNA damage response (Oh et al. 2016). Perhaps, the NTD of Pch2 is required to form a complex with Xrs2 to sustain the *zip1Δ* checkpoint. Alternatively, mutation of the basic motif in Pch2’s NTD may disrupt the AAA+ hexameric complex thus preventing its binding to normal Pch2 chromosomal target sites. Indeed, the ATPase-dead Pch2-K320A mutant version, defective in ATP binding, also fails to form a stable AAA+ complex (Wendler et al. 2012; Chen et al. 2014; Herruzo et al. 2016) and, like Pch2-ntd6A and Pch2-ntd3A, it is unable to localize to either the rDNA or the SC (Herruzo et al. 2016) (Fig. 3c, d). However, ATPase activity *per se* is not required for Pch2 localization because the ATP hydrolysis-deficient Pch2-E399Q version does localize normally to both rDNA and SC despite being inactive, as manifested by its inability to exclude Hop1 from the nucleolar area (Herruzo et al. 2016) (Fig. 2b). Future experiments will address these and other possibilities to delineate the precise role of the essential NTD motif identified in this work.

**Table 2.**
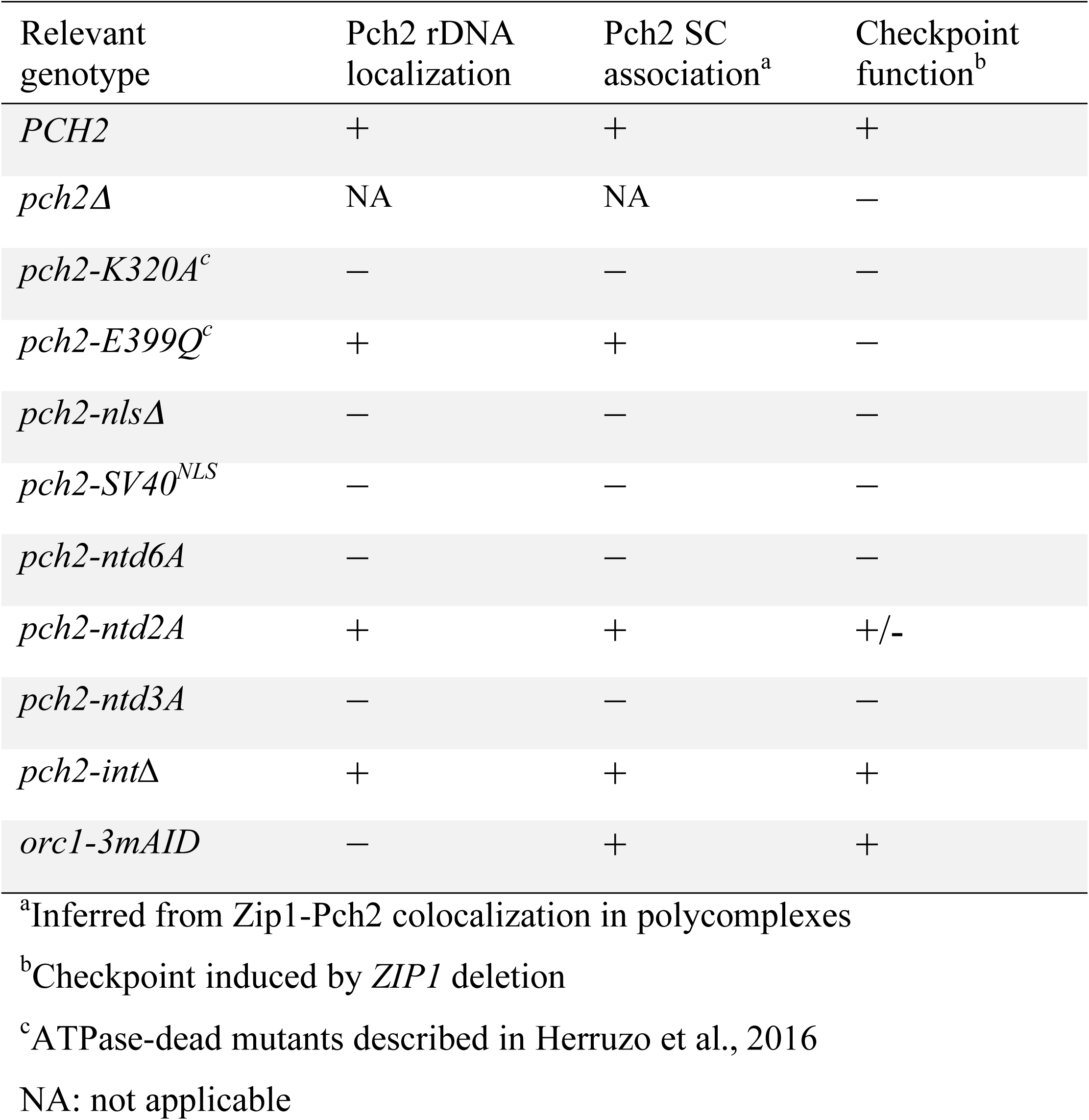
Summary of functional and localization analysis of Pch2

| Relevant genotype | Pch2 rDNA localization | Pch2 SC association<sup>a</sup> | Checkpoint function<sup>b</sup> |
| --- | --- | --- | --- |
| *PCH2* | + | + | + |
| *pch2Δ* | NA | NA | — |
| *pch2-K320A<sup>c</sup>* | — | — | — |
| *pch2-E399Q<sup>c</sup>* | + | + | — |
| *pch2-nlsΔ* | — | — | — |
| *pch2-SV40<sup>NLS</sup>* | — | — | — |
| *pch2-ntd6A* | — | — | — |
| *pch2-ntd2A* | + | + | +/- |
| *pch2-ntd3A* | — | — | — |
| *pch2-intΔ* | + | + | + |
| *orc1-3mAID* | — | + | + |

<sup>a</sup>Inferred from Zip1-Pch2 colocalization in polycomplexes<sup>b</sup>Checkpoint induced by *ZIP1* deletion<sup>c</sup>ATPase-dead mutants described in Herruzo et al., 2016
NA: not applicable

As an additional strategy to elucidate whether the nucleolar Pch2 population is involved in the meiotic recombination checkpoint response, in this work we have also studied the participation of Orc1 in this surveillance mechanism and its requirement for targeting Pch2 to distinct chromosomal locations. Orc1 is an essential component of the Origin Recognition Complex (ORC), which is necessary for initiation of DNA replication during both the mitotic and meiotic cell cycles (Bell et al. 1993; Vader et al. 2011). Besides the replicative function, Orc1 collaborates with Pch2 in maintaining meiotic stability of the rDNA array by preventing DSB formation and the unwanted non-allelic homologous recombination that could potentially arise (Vader et al. 2011). Curiously, Orc1 protein levels are meticulously regulated during meiosis by intricate transcriptional and post-transcriptional mechanisms that ultimately rely on the Ndt80 transcription factor (Xie et al. 2016). Ndt80 is a key target of the meiotic recombination checkpoint raising the possibility of a functional coupling between Orc1 levels and the control of exit from prophase I by the status of checkpoint activation.

In our work, we describe the localization pattern of Orc1 on meiotic chromosomes. In accordance with the multiple replication origins in the rDNA repeats representing ORC binding sites, Orc1 often accumulates on this region partially colocalizing with Pch2. In addition, we also find a general distribution of Orc1 throughout meiotic chromatin likely reflecting Orc1 binding to genomic replication origins. Although we cannot discard a sensitivity issue, Pch2 appears to be absent from these sites suggesting that Orc1 and Pch2 interaction may occur exclusively in the nucleolus. Consistent with this notion, we show that Orc1 is specifically required for localization of Pch2 to the rDNA, but not for its association with SC proteins. Like Pch2, the Orc1 protein also belongs to the AAA+ family of ATPases (Duncker et al. 2009). While Pch2 forms homohexamers in vitro (Chen et al. 2014), the possibility of heteromeric complexes between Pch2 and Orc1 has been suggested (Vader 2015). Supporting this hypothesis, we note that Pch2 nucleolar localization, but not other Orc1 essential functions, is extremely sensitive to Orc1 fusion to small tags that could somehow disrupt proper complex structure. If this were the case, these Pch2-Orc1 heteromeric complexes would be involved exclusively in the rDNA-related functions, whereas Pch2 homohexamers would possess the capacity for removing Hop1 only from synapsed chromosomes. Alternative, it is also possible that nucleolar Pch2 catalytic activity on the Hop1 substrate does not require Orc1, whose main function would be targeting Pch2 to the rDNA to exert the protective function on undesirable DSB formation.

The *orc1-3mAID* allele generated in this work solely compromises Pch2 nucleolar localization allowing us to unequivocally address the direct contribution of this particular genomic location to Pch2’s role in the *zip1Δ*-induced meiotic checkpoint. We show multiple cytological and molecular pieces of evidence demonstrating that in the absence of Orc1 the checkpoint-launching response remains intact, thus indicating that Orc1, and hence nucleolar Pch2, are dispensable for the activation of this quality control mechanism. In the *zip1Δ* mutant lacking the central region of the SC, and thus triggering the checkpoint, Pch2 is only detectable in the rDNA by the chromosome spreading technique. In auxin-treated *zip1Δ orc1-3mAID* nuclei, Pch2 is no longer detected on chromosomes, but the checkpoint is still active. The DNA content profile of the *zip1Δ orc1-3mAID* double mutant is similar to that of *zip1Δ* (Fig. S5e) and, importantly, its strong delay in meiotic progression still relies on Pch2 (Fig. 7b), supporting the conclusion that the meiotic arrest of *zip1Δ orc1-3mAID* results from activation of the Pch2-dependent meiotic recombination checkpoint and not from other unrelated defects. Our results, therefore, open the question of the precise localization and/or distribution of the Pch2 population relevant for the checkpoint response when Zip1 is absent. Although a technical sensitivity issue in Pch2 localization studies cannot be ruled out, it is also possible that a fraction of Pch2 loosely associated to chromatin is responsible for the checkpoint role. In line with this possibility, it has been suggested that the *Drosophila* PCH2 protein exerts its meiotic checkpoint function from a location associated to the nuclear envelope, but at a distance from the chromosomes (Joyce and McKim 2010). Previous studies have revealed a correlation between Pch2 nucleolar mislocalization and meiotic checkpoint deficiency in *sir2* and *dot1* mutants (San-Segundo and Roeder 1999; San-Segundo and Roeder 2000; Ontoso et al. 2013; Cavero et al. 2016). However, we show here that the checkpoint remains intact when Pch2 is removed from the nucleolus upon Orc1 depletion. Therefore, Dot1 and Sir2 may also control the nucleolar-independent population of Pch2 important for checkpoint function. Consistent with the results presented here, the meiotic recombination checkpoint is functional in *zip1Δ rdnΔ* strains lacking the rDNA array on chromosome XII (San-Segundo and Roeder 1999). In this *rdnΔ* scenario, Pch2 shows a substantial redistribution to chromosome ends, and both Pch2 telomeric localization and checkpoint function become dependent on the Sir3 silencing factor, which is not normally required for *zip1Δ* arrest in *RDN*^+^ cells. Curiously, the budding yeast Sir3 protein is a paralog of Orc1 that arose by gene duplication and subsequent functional specialization during evolution (Hanner and Rusche 2017). Thus, multiple regulatory networks impact on Pch2 function and localization in different circumstances. In sum, our results have contributed to narrow down the factors impinging on at least some of the paramount roles of Pch2, such as the *zip1Δ*-induced checkpoint response. Additional future studies will be aimed to discriminate the critical spatiotemporal regulatory mechanisms underlying the meiotic functions of this enigmatic conserved meiotic protein.

## MATERIALS AND METHODS

### Yeast strains and meiotic time courses

The genotypes of yeast strains are listed in Supplementary Table S2. All strains are in the BR1919 or BR2495 background (Rockmill and Roeder 1990). The *zip1Δ∷LEU2, zip1Δ∷LYS2, ndt80Δ∷LEU2, ndt80Δ∷kanMX3, pch2Δ∷URA3* and *pch2Δ∷TRP1* gene deletions were previously described (Herruzo et al. 2016). The *spo11Δ∷natMX4* deletion was generated using a polymerase-chain reaction (PCR)-based approach (Goldstein and McCusker 1999). N-terminal tagging of Pch2 with three copies of the -HA or -MYC epitopes was previously described (San-Segundo and Roeder 1999; Herruzo et al. 2016). The *ORC1-6HA* and *orc1-3mAID* constructs were generated by a PCR-based method using the pYM16 (Janke et al. 2004) and pMK152 (Nishimura and Kanemaki 2014) plasmids, respectively. To direct expression of the *Oryza sativa TIR1* gene during meiosis in yeast for the auxin-induced degron technique, *P*_*HOP1*_*-OsTIR1* was targeted to the genomic *ura3-1* locus by *Stu*I digestion of pSS346 (see below). The *pch2-nlsΔ, pch2-SV40*^*NLS*^ and *pch2-ntd3A* mutations were introduced into the genomic *3HA*-*PCH2* locus using the *delitto perfetto* method that leaves no additional marker (Stuckey et al. 2011). Essentially, the CORE cassette (*URA3-kanMX4*) was first inserted into the *3HA*-*PCH2* gene in the vicinity of the location where the mutation was to be made. Then, the strains carrying *3HA*-*pch2-CORE* were transformed with DNA fragments containing the desired mutation and homologous flanking sequences to both sides of the CORE insertion point to evict the cassette. 5-fluoroorotic acid (FOA)-resistant and G418-sensitive clones were selected and further checked for the presence of the desired mutation. Generation of *pch2-K320A* and *pch2-E399Q* was previously reported (Herruzo et al. 2016). Strains harboring N-terminal tagging of Pch2 with GFP (*GFP-PCH2*) were also constructed using the *delitto perfetto* approach. Basically, a PCR fragment containing the *PCH2* promoter followed by GFP inserted at the second codon of *PCH2* with a five Gly-Ala linker in between (Fig. S3A) was transformed into a strain carrying the CORE cassette close to the 5’ end of *PCH2* and correct FOA-resistant clones were selected. All constructions and mutations were verified by PCR analysis and/or sequencing. The sequences of all primers used in strain construction are available upon request. All strains were made by direct transformation of haploid parents or by genetic crosses always in an isogenic background. Sporulation conditions for meiotic time courses have been described (Ontoso et al. 2013). To score meiotic nuclear divisions, samples were taken at different time points, fixed in 70% Ethanol, washed in phosphate-buffered saline (PBS) and stained with 1 μ g/ μ l 4’,6-diamidino-2-phenylindole (DAPI) for 15 min. At least 300 cells were counted at each time point. Meiotic time courses were repeated several times; averages and error bars from at least three replicates are shown.

### Plasmids

The plasmids used are listed in Supplementary Table S3. The pSS346 plasmid, in which *OsTIR1* is placed under control of the *HOP1* promoter, was constructed by cloning a PCR-amplified 660 bp fragment containing the *HOP1* promoter flanked by *Eco*RI-*Spe*I into the same sites of pMK200 to replace the *ADH1* promoter by the *HOP1* promoter. The different *pch2* mutations in pSS338, pSS358, pSS362, pSS363 and pSS364 were generated following essentially the procedure described in the Q5 site-directed mutagenesis kit (New England Biolabs) using the pSS75 plasmid as template. To analyze the localization of functional GFP-Pch2 in live meiotic prophase cells, the pSS393 plasmid was constructed using several cloning steps. Essentially, pSS393 is a pRS314-derived centromeric plasmid harboring the *HOP1* promoter to drive the prophase-specific expression of the *GFP* coding sequence fused at the second codon of the *PCH2* ORF lacking the intron (Fig. 4b). A flexible linker of five Gly-Ala repeats was also placed between GFP and the Pch2 N-terminus. The pSS396 and pSS397 plasmids driving the production of GFP-Pch2-nlsΔ and GFP-Pch2-SV40^NLS^, respectively, were derived from pSS393 by using the Q5 site-directed mutagenesis procedure (pSS396) or the NEBuilder assembly kit (New England Biolabs) (pSS397). Specific details on plasmid construction and the sequences of all primers used are available upon request.

### Antibody generation

To raise rabbit polyclonal antibodies against Pch2, a DNA fragment encoding amino acids 91–300 was cloned into the pET30a vector (Novagen) for expression in *Escherichia coli*. The His-tagged protein was purified using Ni-NTA resin (Qiagen) following the manufacturer’s instructions and was used for rabbit immunization. Serum was collected after five injections and was affinity purified against the recombinant antigen as described (Petkovic et al. 2005).

To obtain the mouse anti-Hop1 monoclonal antibody, the MonoExpress Gold Antibody service from Genescript was used. In brief, a recombinant fragment of *HOP1* encoding Hop1^20-250^ was used to immunize mice. Hybridomes were generated and positive clones selected for antibody production and affinity purification.

### Western blotting

Total cell extracts were prepared by trichloroacetic acid (TCA) precipitation from 5-ml aliquots of sporulation cultures as previously described (Acosta et al. 2011). The antibodies used are listed in Supplementary Table S4. The ECL, ECL2 or SuperSignal West Femto reagents (ThermoFisher Scientific) were used for detection. The signal was captured on films and/or with a ChemiDoc XRS system (Bio-Rad) and quantified with the Quantity One software (Bio-Rad).

### Cytology

Immunofluorescence of chromosome spreads was performed essentially as described (Rockmill 2009). The antibodies used are listed in Supplementary Table S4. Images of spreads were captured with a Nikon Eclipse 90i fluorescence microscope controlled with MetaMorph software (Molecular Devices) and equipped with a Hammamatsu Orca-AG charge-coupled device (CCD) camera and a PlanApo VC 100×1.4 NA objective. DAPI images were collected using a Leica DMRXA fluorescence microscope equipped with a Hammamatsu Orca-AG CCD camera and a 63x 1.4 NA objective. Images of whole live cells expressing *GFP-PCH2* and *HOP1-mCherry* were captured with an Olympus IX71 fluorescence microscope equipped with a personal DeltaVision system, a CoolSnap HQ2 (Photometrics) camera and 100x UPLSAPO 1.4 NA objective. Stacks of 7 planes at 0.8 μ m intervals were collected. Maximum intensity projections of planes containing Hop1-mCherry signal and single planes of GFP-Pch2 are shown in Fig. 4d and Fig. S4. The linescan tool of the MetaMorph software was used to measure and plot the fluorescence intensity profile across the cytoplasm and nucleus/nucleolus. To determine the nuclear/cytoplasm GFP fluorescence ratio, the ROI manager tool of Fiji software (Schindelin et al. 2012) was used to define the cytoplasm and nuclear (including the nucleolus) areas and the mean intensity values were measured. Background values were subtracted prior to ratio calculation.

### Dityrosine fluorescence assay, sporulation efficiency and spore viability

To examine dityrosine fluorescence as an indicator of the formation of mature asci, patches of cells grown on YPDA plates were replica-plated to sporulation plates overlaid with a nitrocellulose filter (Protran BA85, Whatman). After 3-days incubation at 30°C, fluorescence was visualized by illuminating the open plates from the top with a hand-held 302 nm ultraviolet (UV) lamp. Images were taken using a Gel Doc XR system (Bio-Rad). Sporulation efficiency was quantitated by microscopic examination of asci formation after 3 days on sporulation plates. Both mature and immature asci were scored. At least 300 cells were counted for every strain. Spore viability was assessed by tetrad dissection. At least 144 spores were scored for every strain.

### Statistics

To determine the statistical significance of differences a two-tailed Student *t*-test was used. *P*-Values were calculated with the GraphPad Prism 5.0 software.

## ACKNOWLEDGEMENTS

We are grateful to David Ontoso, Andrés Clemente and Shirleen Roeder for reagents. We also thank Isabel Acosta and Sara González-Arranz for technical assistance, Carlos Vázquez for advice on microscopy analysis, and José Pérez-Martín and Andrés Clemente for helpful discussions and ideas.

## SUPPLEMENTAL DATA

**Fig. S1.**
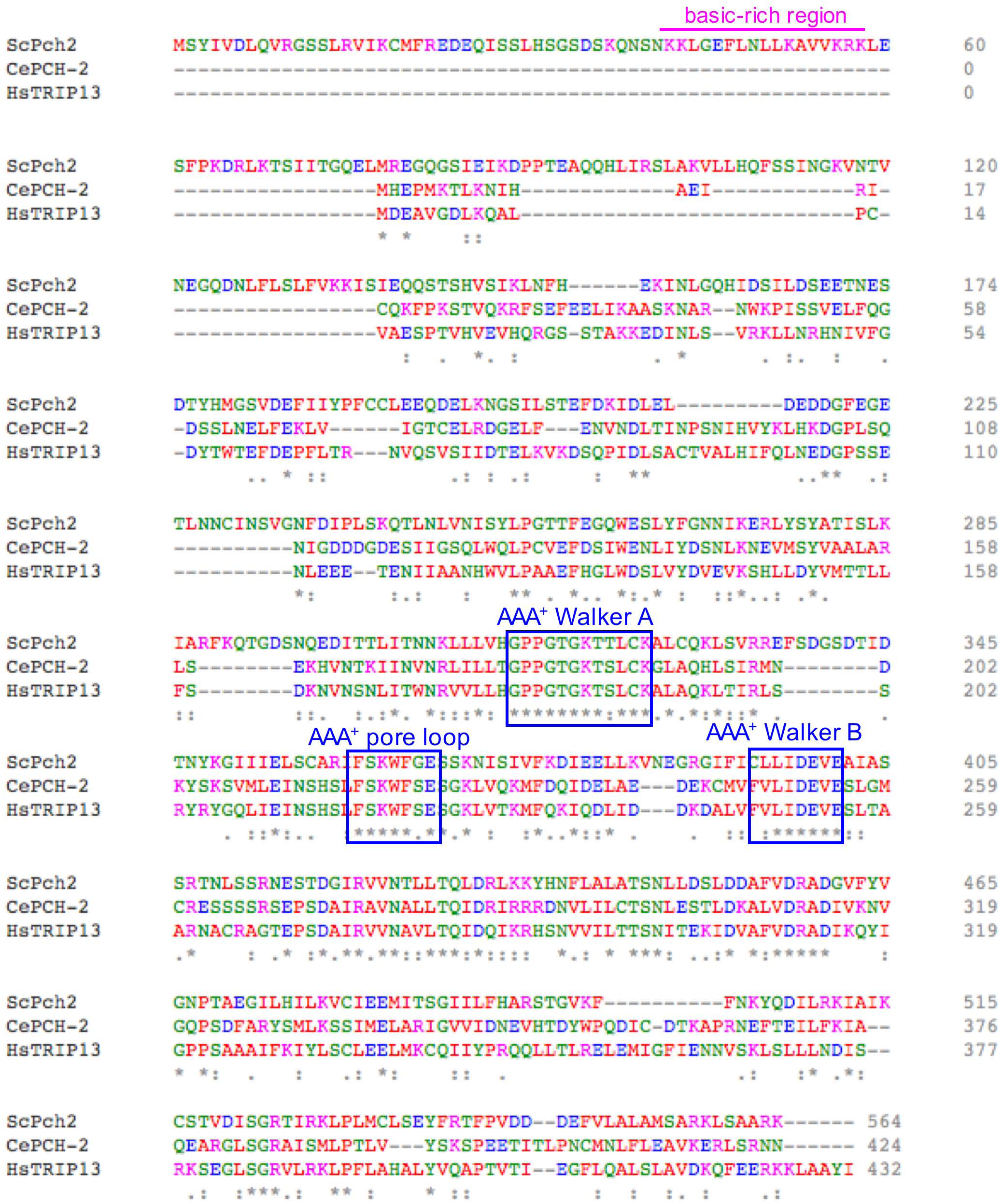
ClustalW alignment of the protein sequences of Pch2 orthologs from *S*. *cerevisiae* (ScPch2), C. *elegans* (CePCH-2) and human (HsTRIP13). The characteristic AAA+ ATPase feahues are boxed (blue). The basic-rich motif in the ScPch2 TD is underlined (magenta). The color code for amino acids is the following: AVFPMILW (small + hydrophobic -Y): red. DE (acidic): blue. RK (basic -H): magenta. STYHC GQ (hydroxyl +sulfhydiyl + amine + G): green. Alignment was performed at https://www.ebi.ac.uk/Tools/msa/clustalo/. |

**Fig. S2.**
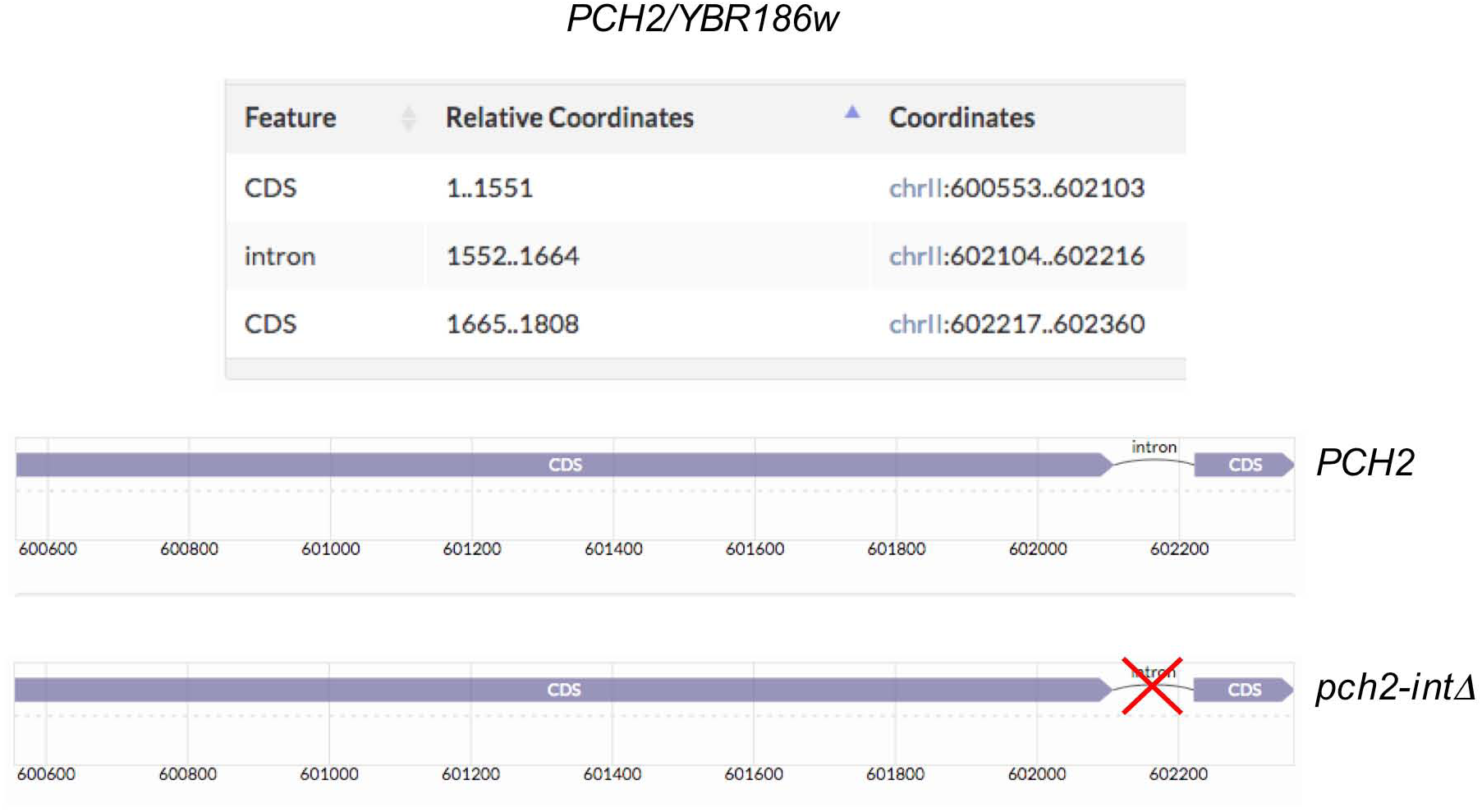
The *S*. *cerevisiae PCH2* locus. Schematic representation of the intron-containing *PCH2* gene displaying the chromosome II coordinates. In the *pch2-int, zip1Δ* allele the intron was deleted, but the coding sequence is unchanged. Data obtained and representation modified from the Saccharomyces Genome Database (https://www.yeastgenome.org/).

**Fig. S3.**
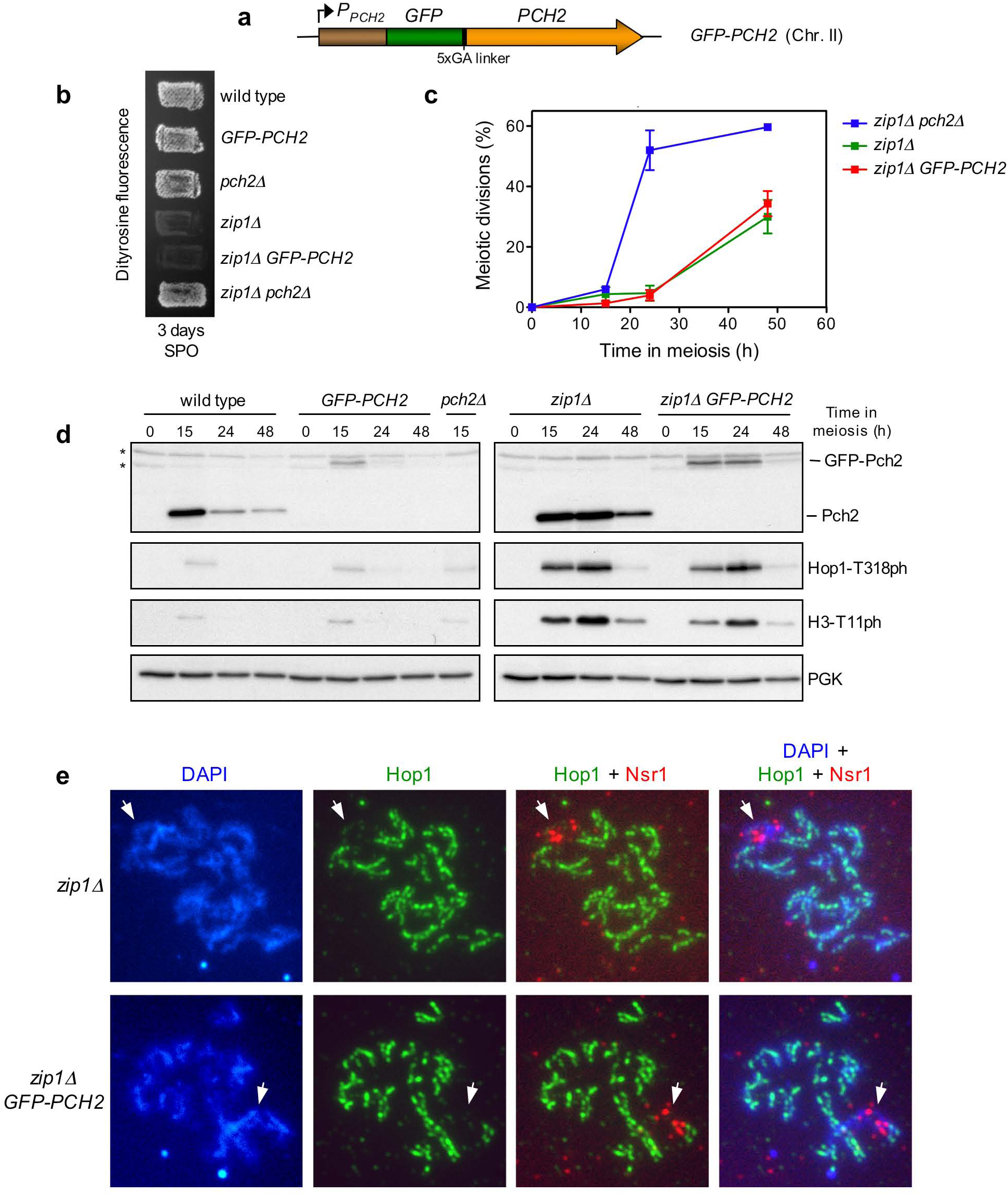
The GFP-Pch2 protein fully supports meiotic checkpoint function. a Schematic representation ofthe *PCH2* gene tagged with *GFP* at its own genomic locus in chromosome II to generate *GFP-PCH2* strains. **b** Dityrosine fluorescence as an indicator of sporulation, was examined after 3 days of sporulation on plates. c Time course analysis of meiotic nuclear divisions; the percentage of cells containing two or more nuclei is represented. Error bars: SD· n=3. d Western blot analysis of Pch2 and GFP-Pch2 production during meiosis (detected with anti-Pch2 antibodies), Hopl-T3l8 phosphorylation and Mekl activation (H3-Tl I phosphorylation). PGK was used as a loading control. e Immunofluorescence of meiotic chromosomes stained with anti-srl antibodies (red), anti-Hop I antibodies (green) and DAPI (blue). Representative nuclei are shown Strains in **(b, c, d, e)** are: DP421 (wild type), DP1508 *(GFP-PCH2),* DP1023 *(pch2Δ),* DP422 *(zip1Δ),* DP1509 *(zip1Δ GFP-PCH2)* and DP1029 *(zip1Δpch2Δ)*.

**Fig. S4.**
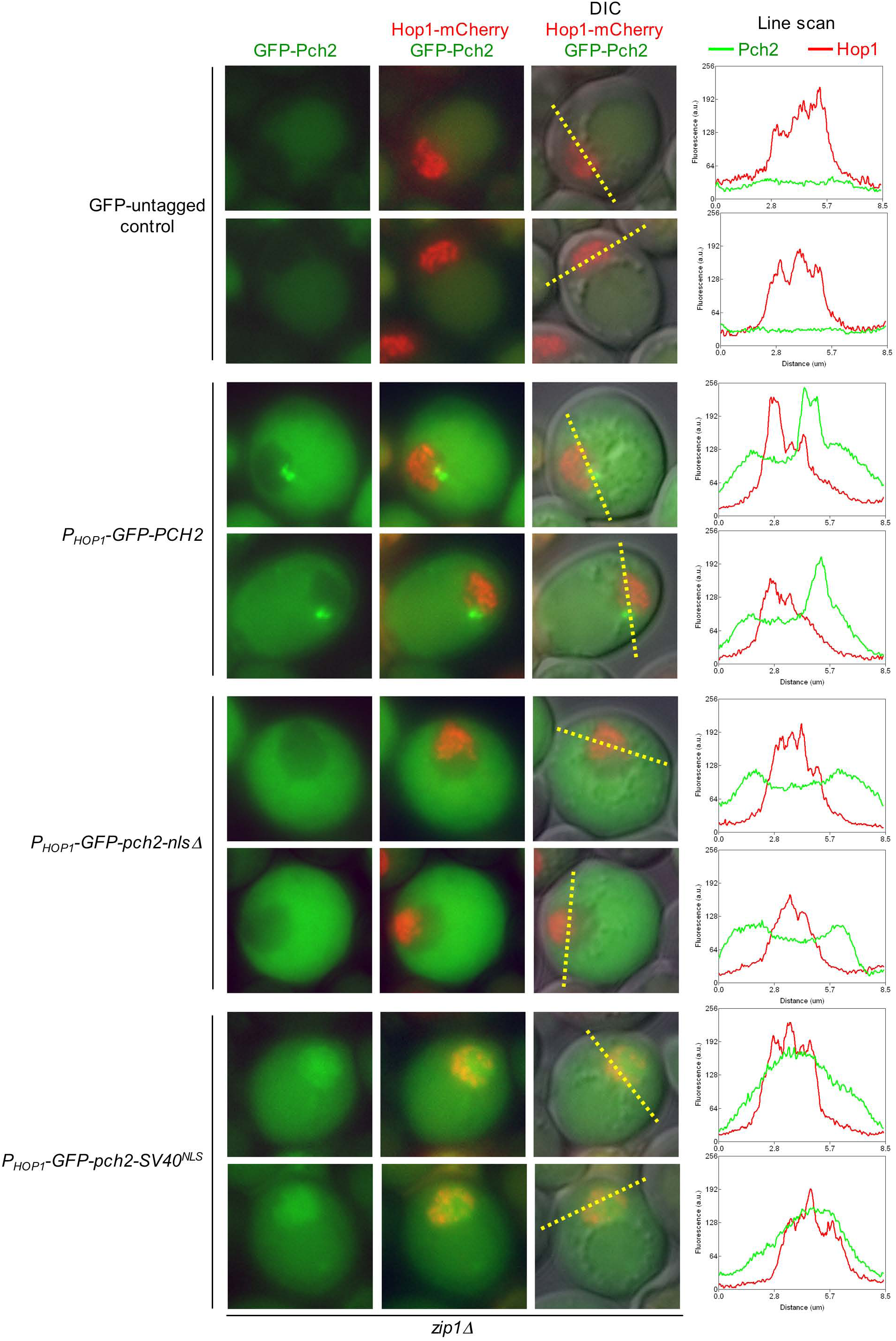
Localization of GFP-Pch2, GFP-Pch2-nlsΔ and GFP-Pch2-SV40NLS in whole meiotic cells. Fluorescence microscopy images, and the corresponding line scan plots, of additional cells to those shown in Fig. 4d. The DP1500 (*zip1Δ*) strain was transformed with pSS393 (*P*_*HoPI*_*GFP-PCH2*), pSS396 (*P*_*HoPI*_*GFP-pch2-nlsΔ*) or pSS397 (*P*_*HoPI*_*GFP-pch2-SV40*^NLS^)

**Fig. S5.**
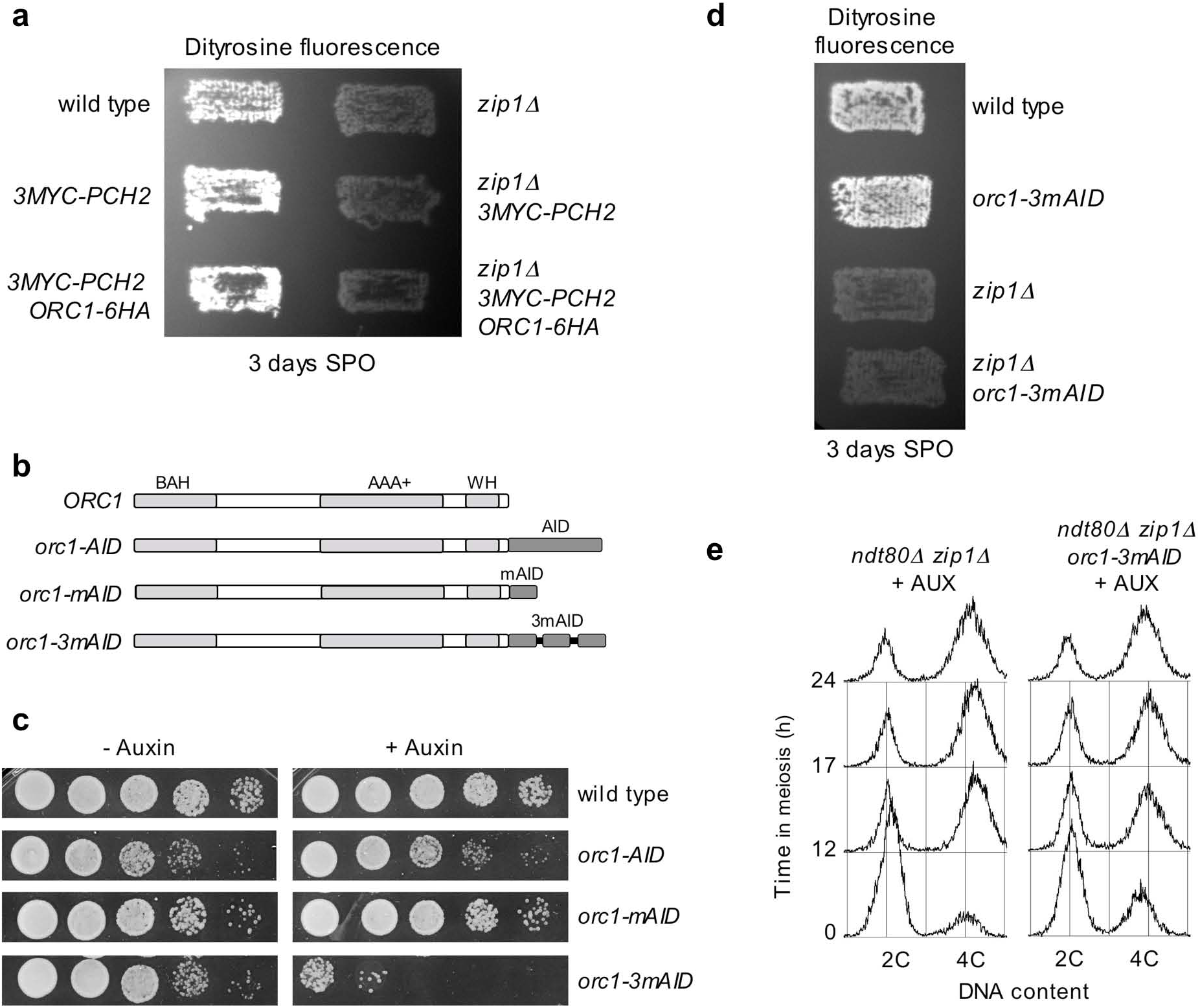
Functional analysis of different tagged versions and degron alleles of *ORC*1. **a** Dityrosine fluorescence, as an indicator ofsporulation, was examined after 3 days of sporulation on plates. Strains in (**a**) are: DP421 (wild type), DP1243 *(3MYC-PCH2),* DP1426 *(3MYC-PCH2 ORCJ-6HA),* DP422 *(zip1Δ*), DPl244 *(zip1Δ 3MYC-PCH2)* and DPl427 *(zip1Δ JMYC-PCH2ORCJ-6HA)*. **b**, **c** Construction and analysis of auxin-induced *orcl* degron alleles. **b** Schematic representation ofthe Ore I protein indicating relevant functional domains (Kawakami et al. 2015). BAH: bromo-adjacent homology. WH: winged helix. The different versions ofthe auxin-induced degron used (AID, mAID and 3mAID) are also depicted. **c** Growth analysis ofthe different *orcl* alleles. Ten-fold serial dilutions oflate log-phase cultures were spotted onto YPDA plates without auxin or containing 500 µM auxin. Representative clones of several transformants tested containing *P*_*ADHI*_*-OsTIRJ* (pMK.200) are shown. **d** Dityrosine fluorescence, as an indicator of sporulation, was examined after 3 days of sporulation on plates. Strains in (**d)** are DP421 (wild type), DP1437 *(orc1-3mAID),* DP422 *(zip1 Δ*) and DP1438 *(zip1 Δ orc1-3mAID)*. ***e*** FACS analysis of DNA content during meiosis. Auxin (500 ***µM)*** was added to cultures 12 h after meiotic induction. Strains in (e) are: DP 1190 *(ndt80 Δ zip1Δ*) and DP1452 *(ndt80 Δzip1 Δ orcl-m3AID)*.

**Fig. S6.**
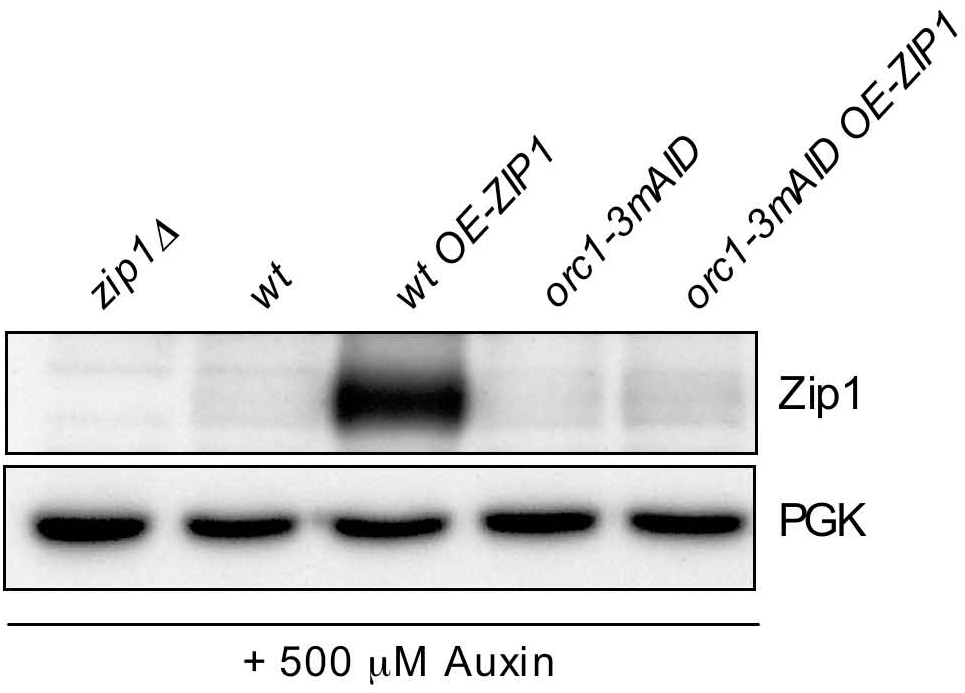
Deficient *ZIP1* overexpression in the *orcl-3mAID* mutant. Western blot analysis of Zipl overproduction. Auxin (500 µM) was added to cultures 12 h after meiotic induction and cell extracts were prepared at 18 hours. Strains are: DP1152 *(zip1Δ*), DP1151 (wt), DP115 I+ pSS343 (wt *OE-ZIP1),* DP1437 *(orcl-3mAID)* and DPl437 + pSS343 *(orc1-3mAID OE-ZIP1)*.

**Fig. S7.**
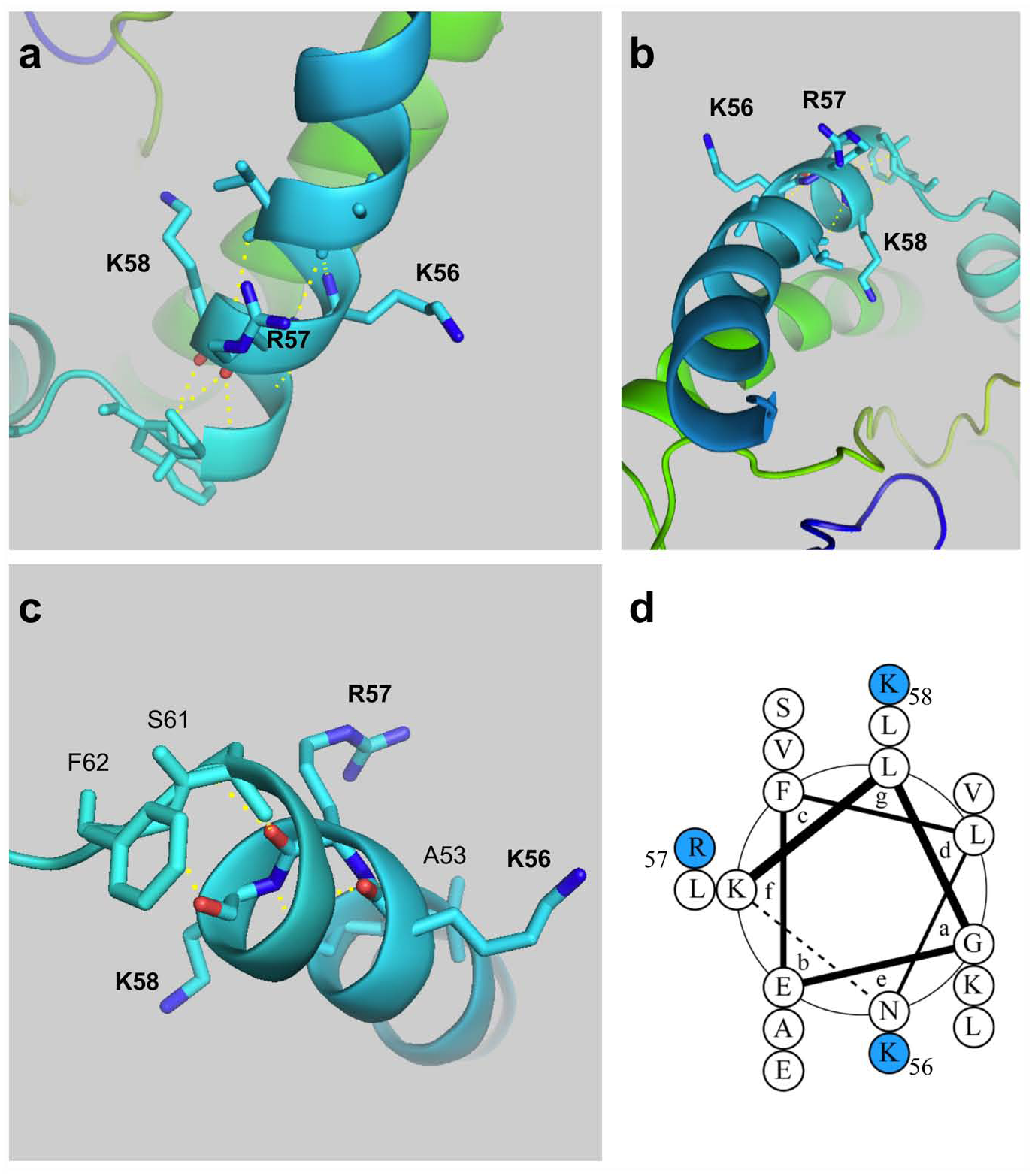
Structural prediction of the basic motif in Pch2’s TD containing the K_56_R_57_K_58_ residues. (a-c) Different views of the positions of the three amino acids K_56_, R_57_, and K_58_ within an α-helix. Relevant amino acids are depicted. Dark blue lines represent basic groups in lysines 56 and 58, and arginine 57. Red lines represent carbonyl groups. Yellow dotted lines represent polar contacts of these three residues with other nearby amino acids. Protein structure was modeled using the Phyre2 web portal (Kelley et al. 2015). Files were visualized, and images obtained, using the PyMOL Molecular Graphics System, Version 2.0 Schrodinger, LLC. **(d)** Two-dimensional distribution of residues **K**_56_**R**_57_**K**_58_ (blue circles) withjn the predicted coiled-coil domain assembled from residues 43-KLGEFL LKAVV<u>KRK</u>LES-61. The diagram was generated using the DrawCoil 1.0 web tool available from https://grigoryanlab.org/drawcoil/.

**Table S1.** Quantitative data corresponding to spread immunolocalization figures

| **Figure 2a** | **Pch2 rDNA localization** |  |  |
| --- | --- | --- | --- |
| **Relevant genotype (Strain)** | **YES (%)** | **NO (%)** | **Nuclei scored** |
| wild type (DP1191) | 100 | 0 | 58 |
| *pch2-nlsΔ* (DP1411) | 0 | 100 | 30 |
| *pch2-SV40<sup>NLS</sup>* (DP1455) | 0 | 100 | 25 |
| *zip1Δ* (DP1190) | 100 | 0 | 20 |
| *zip1Δ pch2-nlsΔ* (DP1412) | 0 | 100 | 31 |
| *zip1Δ pch2-SV40<sup>NLS</sup>* (DP1456) | 0 | 100 | 20 |

| **Figure 2b** | **Pch2 localization at polycomplex** |  |  |
| --- | --- | --- | --- |
| **Relevant genotype (Strain)** | **YES (%)** | **NO (%)** | **Nuclei scored** |
| wild type *OE-ZIP1* (DP1151/pSS343) | 81.8 | 18.2 | 33 |
| *pch2-nlsΔ OE-ZIP1* (DP1408/pSS343) | 0 | 100 | 22 |
| *pch2-K320A OE-ZIP1* (DP1163/pSS343) | 0 | 100 | 20 |
| *pch2-E399Q OE-ZIP1* (DP1287/pSS343) | 78.9 | 21.1 | 19 |
| *pch2-SV40<sup>NLS</sup> OE-ZIP1* (DP1455/pSS343) | 0 | 100 | 21 |

| **Figure 3c** | **Pch2 rDNA localization** |  |  |
| --- | --- | --- | --- |
| **Relevant genotype (Strain)** | **YES (%)** | **NO (%)** | **Nuclei scored** |
| *zip1Δ PCH2* (DP1405/pSS75) | 60 | 40 | 40 |
| *zip1Δ pch2-nlsΔ* (DP1405/pSS338) | 0 | 100 | 40 |
| *zip1Δ pch2-ntd6A* (DP1405/pSS358) | 2.5 | 97.5 | 40 |
| *zip1Δ pch2-ntd2A* (DP1405/pSS363) | 53.7 | 46.3 | 41 |
| *zip1Δ pch2-ntd3A* (DP1405/pSS364) | 0 | 100 | 40 |
| *zip1Δ pch2-intΔ* (DP1405/pSS362) | 62.2 | 37.8 | 37 |
| **Hop1 localization pattern** |  |  |  |
| **Relevant genotype (Strain)** | **Linear (%)** | **Fragmented (%)** | **Nuclei scored** |
| *zip1Δ PCH2* (DP1405/pSS75) | 58 | 42 | 50 |
| *zip1Δ pch2-nlsΔ* (DP1405/pSS338) | 0 | 100 | 40 |
| *zip1Δ pch2-ntd6A* (DP1405/pSS358) | 2.1 | 97.9 | 47 |
| *zip1Δ pch2-ntd2A* (DP1405/pSS363) | 52.9 | 47.1 | 51 |
| *zip1Δ pch2-ntd3A* (DP1405/pSS364) | 2.3 | 97.7 | 43 |
| *zip1Δ pch2-intΔ* (DP1405/pSS362) | 52.5 | 47.5 | 40 |

| **Figure 3d** | **Pch2 localization at polycomplex** |  |  |
| --- | --- | --- | --- |
| **Relevant genotype (Strain)** | **YES (%)** | **NO (%)** | **Nuclei scored** |
| *PCH2* (DP186/pSS75+pSS343) | 68.4 | 31.6 | 19 |
| *pch2-ntd6A* (DP186/pSS358+pSS343) | 0 | 100 | 18 |
| *pch2-ntd2A* (DP186/pSS363+pSS343) | 30.8 | 69.2 | 26 |
| *pch2-ntd3A* (DP186/pSS364+pSS343) | 0 | 100 | 16 |
| *pch2-intΔ* (DP186/pSS362+pSS343) | 57.9 | 42.1 | 19 |

**Figure 5a**
|  | Pch2 rDNA accumulation |  |  |
| --- | --- | --- | --- |
| Relevant genotype (Strain) | YES (%) | NO (%) | Nuclei scored |
| 3MYC-PCH2 ORC1-6HA (DP1426) | 84.7 | 15.3 | 72 |
| *zip1Δ* 3MYC-PCH2 ORC1-6HA (DP1427) | 58.8 | 41.2 | 51 |
| 3MYC-PCH2 (DP1243) | 100 | 0 | 12 |
| *zip1Δ* 3MYC-PCH2 (DP1244) | 100 | 0 | 13 |
|  | Pch2-Orc1 colocalization in the rDNA |  |  |
| Relevant genotype (Strain) | YES (%) | NO (%) | Nuclei scored |
| 3MYC-PCH2 ORC1-6HA (DP1426) | 100 | 0 | 61 |
| *zip1Δ* 3MYC-PCH2 ORC1-6HA (DP1427) | 100 | 0 | 30 |

**Figure 5b**
|  | Orc1 rDNA localization |  |  |
| --- | --- | --- | --- |
| Relevant genotype (Strain) | YES (%) | NO (%) | Nuclei scored |
| 3MYC-PCH2 ORC1-6HA (DP1426) | 100 | 0 | 26 |
|  | Orc1 rDNA accumulation |  |  |
| Relevant genotype (Strain) | YES (%) | NO (%) | Nuclei scored |
| 3MYC-PCH2 ORC1-6HA (DP1426) | 53.9 | 46.1 | 26 |

**Figure 6b**
|  | Pch2 rDNA localization |  |  |
| --- | --- | --- | --- |
| Relevant genotype (Strain) | YES (%) | NO (%) | Nuclei scored |
| wild type + Auxin (DP1151) | 100 | 0 | 16 |
| *orc1-3mAID* + EtOH (DP1437) | 0 | 100 | 15 |
| *orc1-3mAID* + Auxin (DP1437) | 0 | 100 | 18 |
| *zip1Δ* + Auxin (DP1152) | 100 | 0 | 20 |
| *zip1Δ orc1-3mAID* + EtOH (DP1438) | 0 | 100 | 20 |
| *zip1Δ orc1-3mAID* + Auxin (DP1438) | 0 | 100 | 20 |

**Figure 6c**
|  | Hop1 exclusion from the nucleolus |  |  |
| --- | --- | --- | --- |
| Relevant genotype (Strain) | YES (%) | NO (%) | Nuclei scored |
| wild type + Auxin (DP1151) | 100 | 0 | 29 |
| *orc1-3mAID* + Auxin (DP1437) | 0 | 100 | 20 |
| *zip1Δ* + Auxin (DP1152) | 100 | 0 | 23 |
| *zip1Δ orc1-3mAID* + Auxin (DP1438) | 0 | 100 | 21 |

**Figure 6d**
|  | Pch2 localization at polycomplex |  |  |
| --- | --- | --- | --- |
| Relevant genotype (Strain) | YES (%) | NO (%) | Nuclei scored |
| *spo11Δ* + Auxin (DP1425) | 54.2 | 45.8 | 24 |
| *spo11Δ orc1-3mAID* + Auxin (DP1444) | 76.0 | 24.0 | 25 |

**Table S2.**
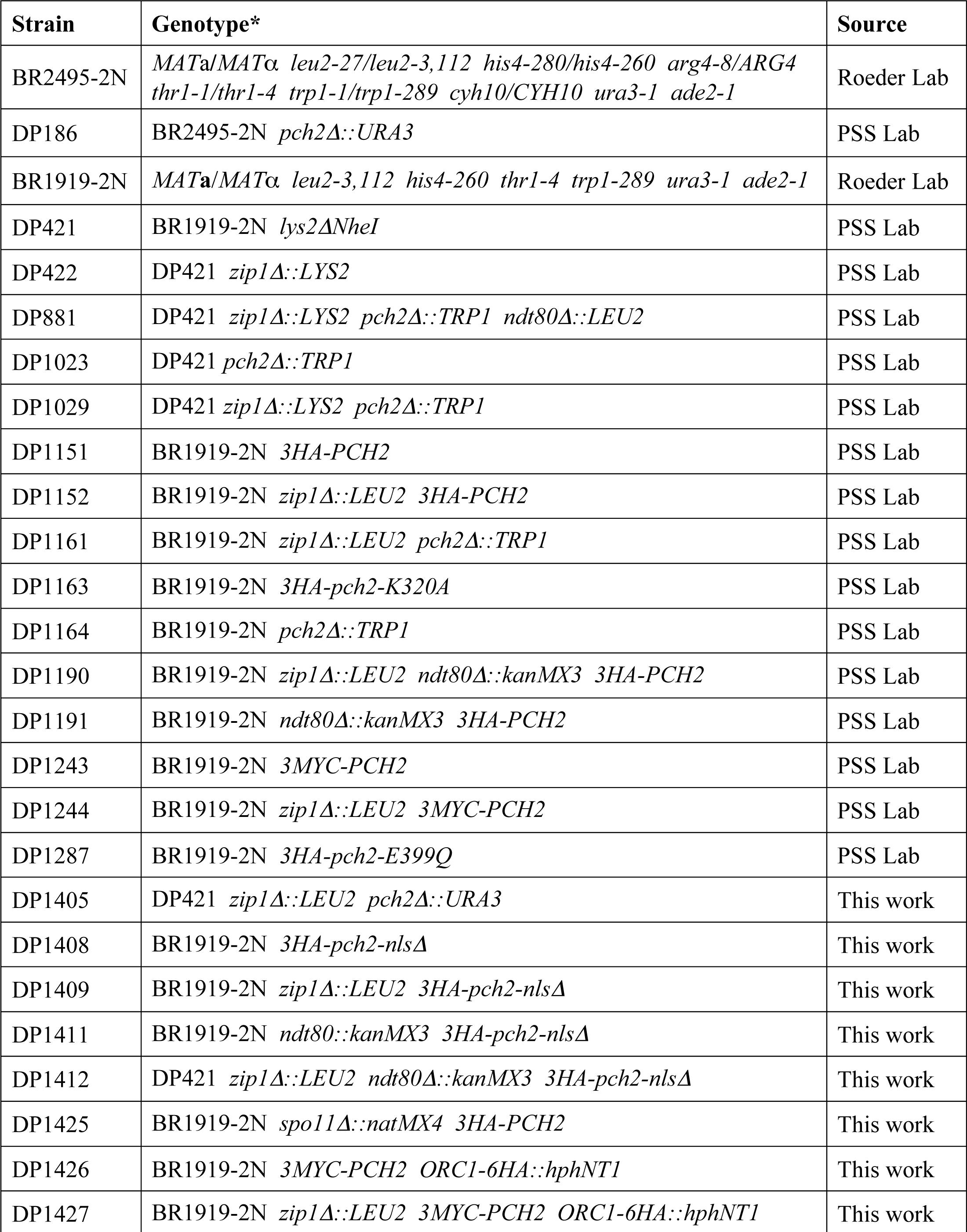
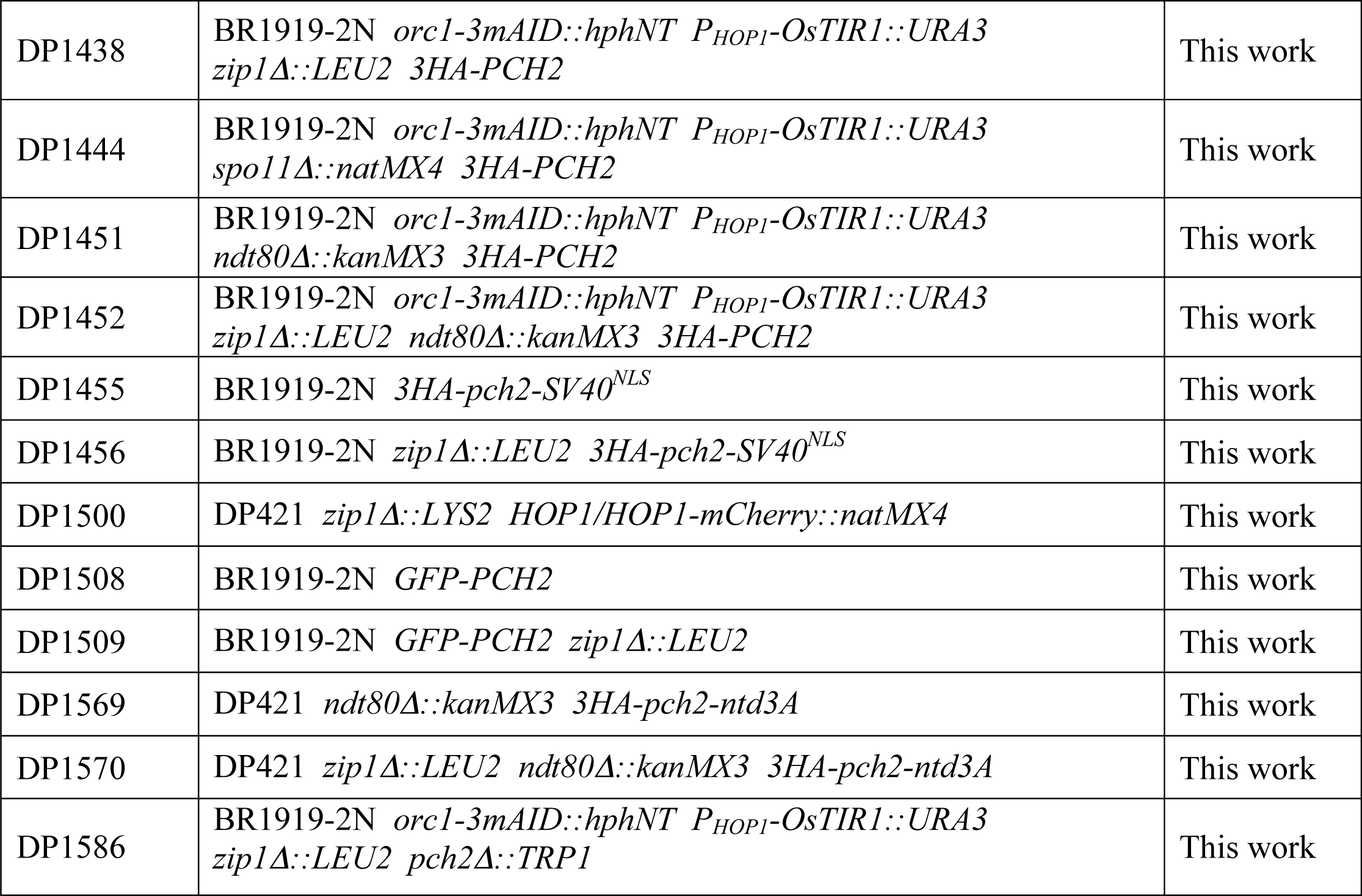
*Saccharomyces cerevisiae* strains.

**Table S3.**
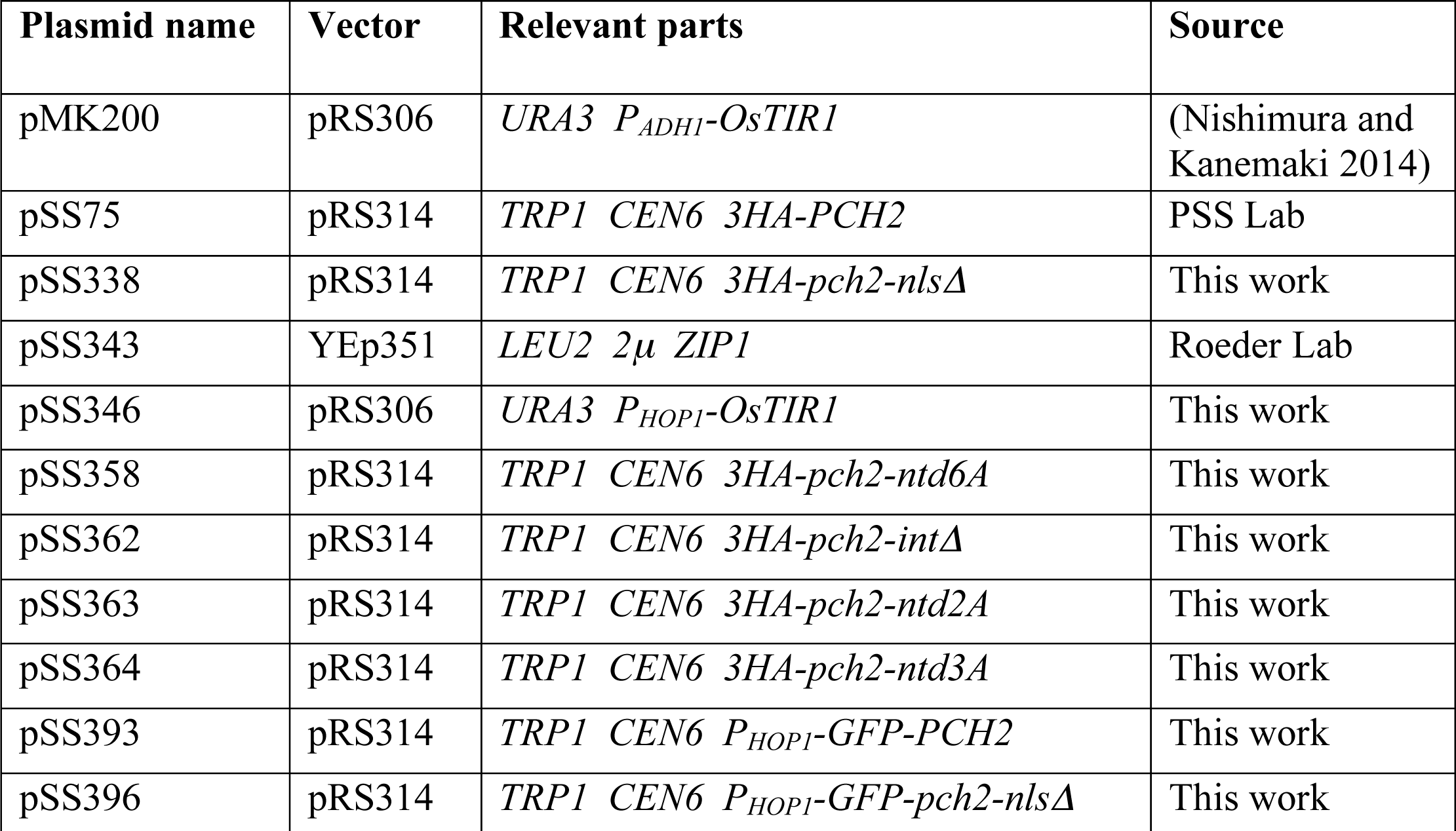
Plasmids.

**Table S4.** Primary antibodies.

| **Antibody** | **Host and type** | **Application\***<br>**(Dilution)** | **Source / Reference** |
| --- | --- | --- | --- |
| Hop1 (5C12E8) | Mouse monoclonal | WB (1:2000) | This work |
| Hop1 | Rabbit polyclonal | IF (1:400) | (Smith and Roeder 1997) |
| Hop1-T318-P | Rabbit polyclonal | WB (1:1000) | (Penedos, *et al.* 2015) |
| H3-T11-P | Rabbit polyclonal | WB (1:2000) | Abcam<br>ab5168 |
| PGK (22C5D8) | Mouse monoclonal | WB (1:5000) | Molecular Probes<br>459250 |
| Pch2 | Rabbit polyclonal | WB (1:2000)<br>IF (1:325) | This work |
| Nsr1 (31C4) | Mouse monoclonal | IF (1:150) | ThermoFisher<br>MA1-10030 |
| Nsr1 (2.3 b) | Mouse monoclonal | IF (1:20) | M. Snyder |
| HA (12CA5) | Mouse monoclonal | WB (1:2000)<br>IF (1:200) | Roche<br>11 666 606 001 |
| Zip1 | Rabbit polyclonal | WB (1:2000)<br>IF (1:200) | (Sym, *et al.* 1993) |
| mAID (1E4) | Mouse monoclonal | WB (1:400) | MBL<br>M214-3 |
| H4-K16ac | Rabbit polyclonal | IF (1:200) | Millipore<br>07-329 |

\*WB, western blot; IF, immunofluorescence

